# Age-specific differences in the dynamics of protective immunity to influenza

**DOI:** 10.1101/330720

**Authors:** Sylvia Ranjeva, Rahul Subramanian, Vicky J. Fang, Gabriel M. Leung, Dennis K. M. Ip, Ranawaka A. P. M. Perera, J. S. Malik Peiris, Benjamin J. Cowling, Sarah Cobey

## Abstract

Influenza A viruses evolve rapidly to escape host immunity, such that individuals can be infected multiple times with the same subtype. The form and duration of protective immunity after each influenza infection are poorly understood. Here, we quantify the dynamics of protective immunity against influenza A virus infections by fitting individual-level mechanistic models to longitudinal serology from children and adults in a household cohort study. We find that most protection in children is explained by antibody titers measured by the hemagglutination inhibition (HI) assay. In contrast, in adults, HI antibody titers explain a smaller fraction of protection. Protection against circulating strains wanes to approximately 50% of peak levels 3.5-7 years after infection in both age groups, and wanes faster against influenza A(H3N2) than A(H1N1)pdm09. Protection against H3N2 lasts longer in adults than in children. Our results suggest that the focus of influenza antibody responses changes over time from the highly mutable hemagglutinin head to other epitopes, consistent with the immunological theory of original antigenic sin, and that this change of focus might affect protection. Additionally, we estimate that imprinting, or primary infection with a subtype of one phylogenetic group, has little to no effect on the risk of non-medically attended influenza infections in adults. We also find no evidence of long-term cross-protection between subtypes. This work underscores the need for longitudinal data on multiple components of the immune response to better understand the development of immunity and differences in susceptibility within populations.

---

Like many antigenically variable pathogens, influenza viruses continuously evolve to escape host immunity. As a consequence, they cause frequent epidemics and infect people repeatedly during their lives. The details of these processes—which are vital to influenza epidemiology, evolution, and the design of effective vaccines—have nonetheless remained surprisingly difficult to pin down despite nearly 70 years of study.

A major challenge is uncertainty about the nature of acquired immunity. Antibodies are the primary means of protection against influenza and impose strong selection on its surface proteins [1, 2]. Antibody responses to influenza are highly cross-reactive, in that antibodies induced by infection or vaccination with one strain often protect against infections with related strains [3, 4]. The hierarchical nature of this cross-reactivity, in which memory responses to conserved antigens tend to dominate over responses to new epitopes, is known as original antigenic sin [5, 6, 7]. It might underlie the phenomenon of imprinting, in which primary infection with one influenza subtype protects against severe disease and death with other subtypes that are closely related phylogenetically [8]. But the specificity and duration of protection after seasonal influenza infection have been difficult to estimate, partly because the relationship between antibody titer and protection appears complex, and also because longitudinal observations of antibody titers and infections are rare. The most common measure of anti-influenza antibody is the hemagglutination inhibition (HI) assay, and HI antibody titers are an established correlate of protection [9]. The HI titer corresponding to 50% protection against infection, commonly cited as 40 [10, 11], may vary by influenza A subtype and host age [12, 13], although measurement error, long intervals between titer measurements, and variable titer changes after infection complicate inferences. Recent models have made progress by incorporating measurement error [14, 15], representing infections as latent states [14, 16, 17], and using titers to historic strains to measure the intervals between infections [14], attack rates [15, 16], and the breadth of the response over time [14, 17]. But the relatively short periods of observation in these studies have made it difficult to estimate some basic quantities in the response to infection, namely, how long protection lasts, and whether antibody titers adequately reflect the strength of protection against infection in individuals over time.

Longitudinal cohorts provide relatively unobscured observations of the dynamics of infection and protection, and mechanistic models allow hypotheses about these dynamics to be tested. We fitted stochastic mechanistic models to influenza antibody titers collected over five years from a large household cohort study including children and adults. These models account for pre-existing immunity, variation in the response to infection, and the possibility that the HI titer is not a good correlate of protection from infection. Their flexibility allows many previous assumptions to be relaxed. For both influenza A subtypes, we estimated the duration of within-subtype and cross-subtype protection, the relationship between HI titer and protection, and the effect of earlychildhood influenza exposures on infection risk later in life. The dynamics inferred from these individual-level models are remarkably consistent with the epidemiological dynamics of the larger population, and they also support immunological theory of how the antibody response to influenza changes with age.

## Results

### Homosubtypic protection correlates better with anti-HA antibodies in children than adults

We fitted models to data from a cohort of 592 adults (> 15 y) and 114 children (≤15 y) followed from 2009 to 2014 in Hong Kong. Members of this cohort were part of a larger household study [18, 19] and were selected because they were not vaccinated for the study and reported no vaccination during the five years of follow-up. The cohort included 337 households with a median size of 2 members (Fig. 4b). Sera were obtained every six months and tested for antibodies to circulating strains of influenza A(H3N2) and A(H1N1)pdm09 via the HI assay.

Antibodies measured by the HI assay are an established correlate of protection for influenza virus infection [10, 20, 21]. Neutralizing antibodies against the dominant surface proteins, hemagglutinin (HA) and neuraminidase (NA), can target different sites on them, and the specificity of the antibody response appears to change with immune history and age [22, 23, 24, 25, 26]. HI assays measure antibodies to HA but not NA, and they disproportionately measure anti-HA antibodies that attach near the receptor binding site toward the top of the HA globular domain. To characterize the role of these antibodies in protection, we tested a simple hypothesis about the dynamics of susceptibility after infection: protection from infection could be associated with HI titer, the time since last infection (a potential correlate of other antibody and broader immune responses), or a mixture of the two. We define an individual’s susceptibility to a subtype as the probability of infection given exposure. Rewriting the hypotheses mathematically, we propose the susceptibility of an individual *i* to subtype *s* at time *t*, *q*_*i*,*s*_ (*t*), is a function of
1. *HI-correlated factors.* An individual’s susceptibility can be measured by the HI titer to a representative circulating strain. We assume that HI-correlated susceptibility, *q*_1*i*,*s*_ (*t*), is a logistic function of the current titer [10, 11], with the shape of the curve set by the titer at which 50% of subjects are protected from infection (Fig. 1, Step 1b). This 50% protective titer is defined for each age group *a ∈* {child, adult}, *TP50_ai_*_,*s*_ (Eq. 9),

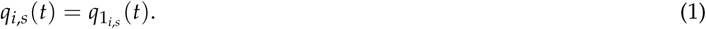
2. *Non-HI-correlated factors.* Susceptibility can be explained by the time since last infection with that subtype (Fig. 1, Step 1b). Susceptibility determined by non-HI-correlated protection, *q*_2*i*,*s*_ (*t*), is a function that starts at 0 (no susceptibility) immediately after infection. The susceptibility increases as protection wanes exponentially at rate *w*_nonspecific,*ai*, *s*_ (Eq. 10),

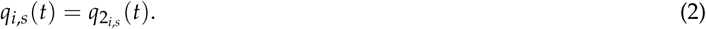

**Figure 1:**
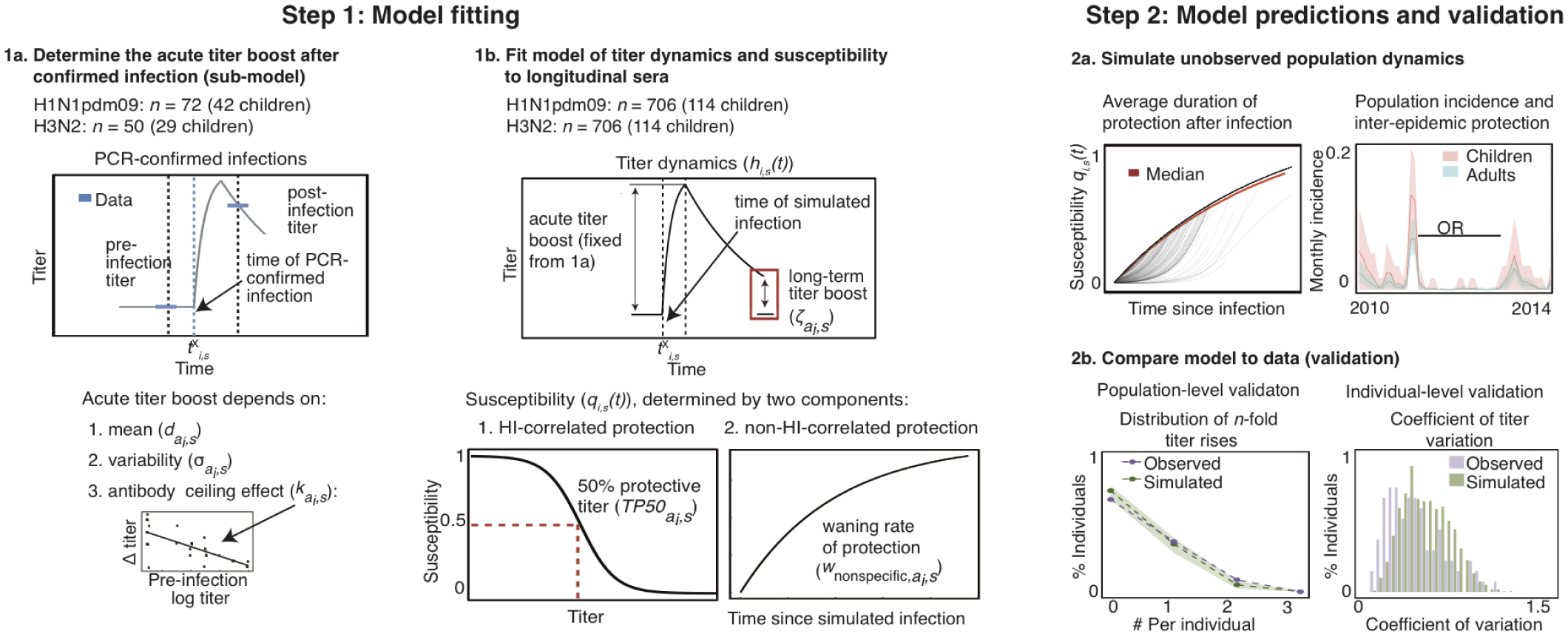
Schematic of modeling approach. Steps 1 and 2 are performed for each subtype. **Step 1**: (**a**) First, the sub-model of short-term post-infection titer dynamics is fitted to a subset of the data. This subset includes the time of PCR-confirmed infection and the immediate pre- and post-infection titers. The mean and standard deviation of the acute titer boost and the mean’s dependence on pre-infection titer (the antibody ceiling effect) are fitted. (**b**) Next, fixing the parameters associated with short-term titer dynamics (Step 1a), the full model is fitted to titers from the entire cohort. The contribution of HI-correlated and non-HI-correlated protection, the titer waning rate, the 50% protective titer, and the long-term boost after infection are estimated. **Step 2**: (**a**) The duration of protection and inter-epidemic protection are estimated from simulating population-level dynamics from the best-fit model in Step 1b. From the latent infections and susceptibility for each individual, we track the loss of protection after infection. We also estimate the incidence and the odds ratios (OR) of protection between epidemics. (**b**) Simulation enables additional checks of the model. We compare the simulated and observed distributions of *n*-fold titer rises and coefficients of titer variation among individuals.

Titers in this model are still informative as indicators of infections, but they do not affect infection risk. We evaluate the contribution of each component by fitting a weighted susceptibility model,

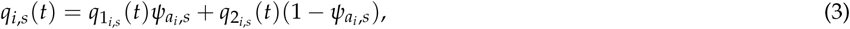

where *ψ*_*ai*,*s*_ measures the contribution of HI-correlated protection to susceptibility in children (*a*_*i*_ = children) and adults (*a*_*i*_ = adults). The value of *ψ*_*ai*,*s*_ therefore distinguishes between models in which protection is completely HI-correlated (*ψ*_*ai*,*s*_ = 1), models in which protection is completely non-HI-correlated (*ψ*_*ai*,*s*_ = 0), and models in which protection is predicted by a combination of the two components (0 < *ψ*_*ai*,*s*_ < 1). To estimate the contribution of HA-head-directed antibodies to protection from influenza infection in children and adults, Eq. 3 was incorporated into a partially observed Markov model that simulates individuals’ latent (unobserved) HI titers and susceptibility to infection over time while simultaneously accounting for measurement error (Fig. 1, Step 1b; Methods). The model assumes that infection can change the antibody titer, which allows infection events and thereby latent susceptibility (*q*_*i*,*s*_ (*t*)) to be inferred from longitudinal sera.

In the model, infection acutely boosts an individual’s titer, which then wanes slowly over one year, potentially leaving a long-term boost that does not wane. To increase accuracy in modeling these acute boosts, we took advantage of 112 PCR-confirmed infections and pre- and post-infection titers from this study to fit the mean and standard deviation of the titer rises (Fig. 1, Step 1a; section S1). The acute boost was higher for H3N2 than for H1N1pdm09 in both children and adults, but there was no significant difference in boost sizes by age in either subtype (Table S1). We found evidence of an antibody ceiling effect, whereby individuals with higher pre-infection titers have smaller boosts (section S1.2). After fitting this “sub-model” to describe the relationship between infection and short-term titer changes, we then fixed its parameters to fit the full model of titer dynamics to all 706 individuals. For children and adults, the full model estimates the contribution of HI-correlated and non-HI-correlated factors to protection (Eq. 3), the magnitude of the long-term titer boost, the 50% protective titers (for Eqs. 1 and 3), and the rate of waning of non-HI-correlated protection (for Eqs. 2 and 3) (Fig. 1, Step 1b). Additionally, because some subjects in the study belong to the same household, we estimate the contribution of infected household members to an individual’s force of infection, relative to that of the community. Simulating from the maximum likelihood estimates of the best model yields additional information, including the typical duration of protection after infection, attack rates in different epidemics, and the odds ratios of infection from one epidemic to the next (Fig. 1, Step 2a). These simulations are also useful for checking how well the model reproduces different features of the data (Fig. 1, Step 2b).

For both subtypes, protection in children is HI-correlated (*ψ*_children,H3N2_ = 1.0, 95% CI: (0.8, 1.0); *ψ*_children,H1N1pdm09_ = 1.0, 95% CI: (0.8, 1.0); Table 1), whereas in adults, time since infection better predicts protection (*ψ*_adults,H3N2_ = 0.0, 95% CI (0.0, 0.2); *ψ*_adults,H1N1pdm09_ = 0.1, 95% CI (0.07, 0.12)). This result suggests that early in life, protection against influenza virus infection is dominated by immune responses that correlate well with HI titer, such as antibodies to the top of the HA head. However, over time, other immune responses dominate, such that time since infection becomes a better predictor of protection than HI titer. This result is consistent with the observation that more children than adults in this study have detectable baseline HI titers, and children have higher mean baseline HI titers, to circulating strains (Fig. S1, Methods).

**Table 1:**
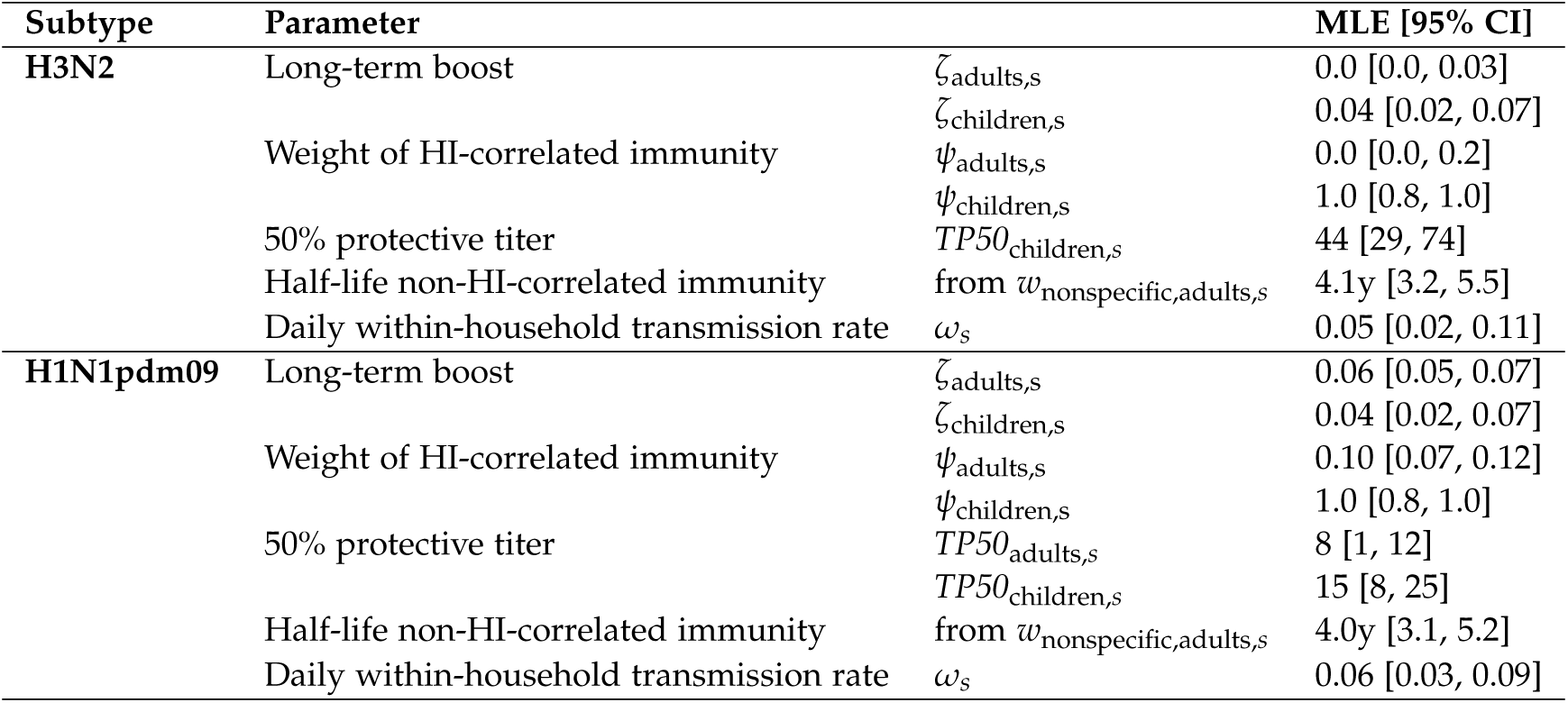
Maximum likelihood estimates and 95% confidence intervals (CI).

To estimate the duration of protection in adults, we simulated from the fitted models for each subtype. Using 1000 replicate simulations, we tracked the latent susceptibility after infection. For each individual at any time, this susceptibility is given by the weighted average of two components, one set by the titer and the other by the time since infection (Eq. 3). In adults, the model favors non-HI-correlated protection against H3N2, with a half-life of 4.1 y (95% CI (3.2, 5.5)) (Table 1). In contrast to H3N2, however, HI-correlated protection contributes slightly to adults’ protection against H1N1pdm09 (*ψ*_adults,H1N1pdm09_ = 0.1 (95% CI (0.07, 0.12)). The associated low 50% protective titer (*TP*_50,adults,H1N1pdm09_ = 8 (95% CI (1,12), Fig. S2) implies that most individuals (i.e., even individuals with titer <10, the lower limit of detection by HI assays) benefit from minor additional protection after infection. When we plot the latent susceptibility after each infection with each subtype, we see that the individual trajectories in Fig. 2A and B follow the curve describing the change in non-HI-correlated protection (Eq. 10). The contribution of HI-correlated protection to the post-infection protection dynamics of H1N1pdm09 creates mild individual-level variability arising from differences in pre-infection titers. In aggregate, adults’ susceptibility to H1N1pdm09 thus wanes with a half-life of approximately 6.4 y (95% quantile: (4.8, 6.7)), reflecting a significantly slower loss of protection than for H3N2. Infection in adults produces only a small durable titer boost to H1N1pdm09 (*ζ*_adults,H1N1pdm09_ = 0.06 (95% CI: 0.05, 0.07)) and a negligible durable boost to H3N2 (*ζ*_adults,H3N2_ = 0.0, (95% CI: 0.0, 0.03); Table 1).

**Figure 2:**
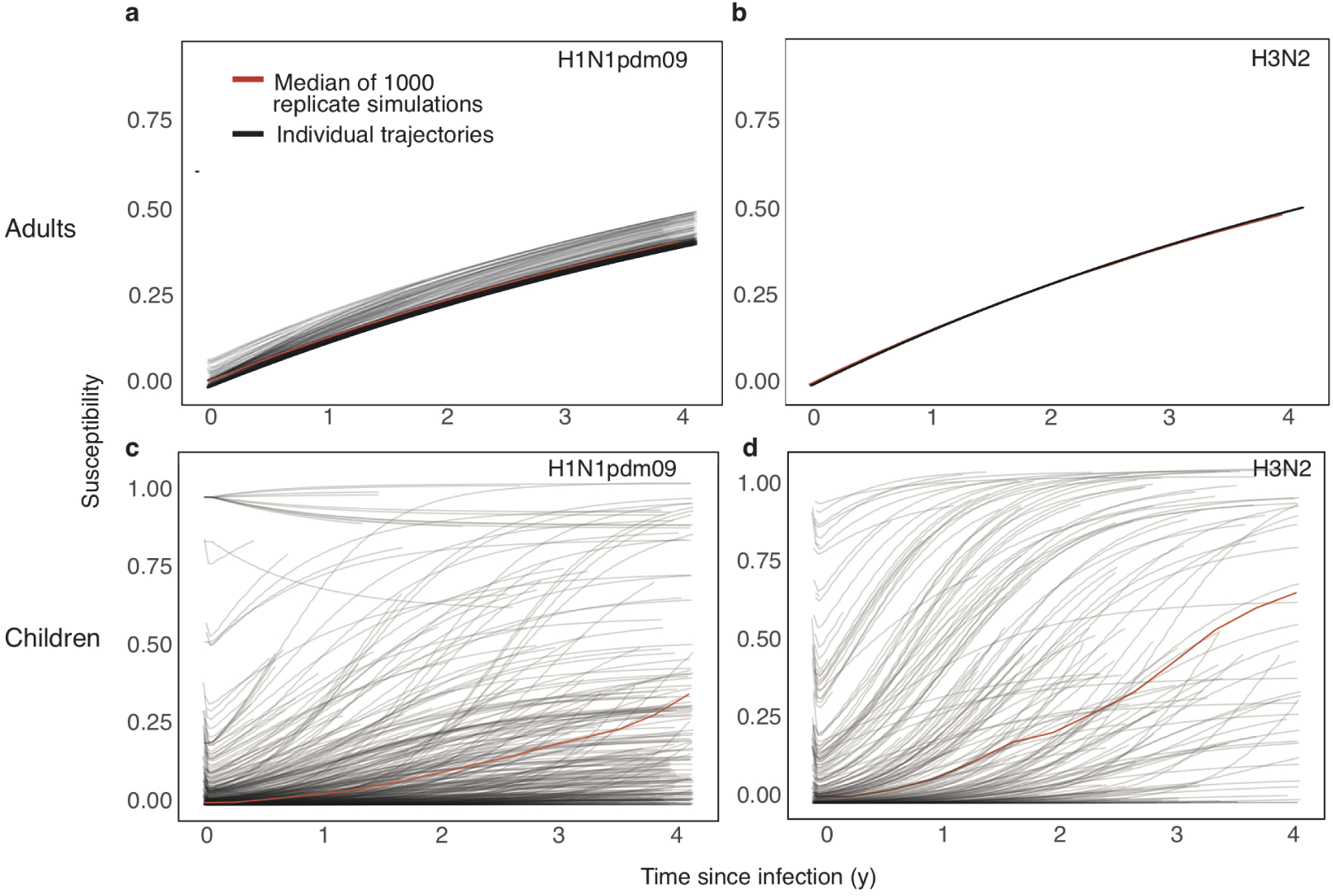
Susceptibility after simulated infection (at time *t* = 0) for adults with H1N1pdm09 (**a**) and H3N2 (**b**) and for children with H1N1pdm09 (**c**) and H3N2 (**d**). The black lines represent individual trajectories from one simulation, and the red line represents the median among individuals over 1000 replicate simulations. Curves from individual trajectories are truncated at points corresponding to the end of the study.

Compared to adults, children have a more variable duration of protection. Because susceptibility in children depends only on HI titer, the dynamics of individual protection are sensitive to pre-infection titers and differences in the magnitude of the acute boost post-infection. For both H1N1pdm09 and H3N2, we estimated substantial variation in the short-term titer dynamics after PCR-confirmed infection (section S1.2). The variability arises both from stochastic variation in the magnitude of the short-term titer boost and from the antibody ceiling effect (Table S1). Protection in children wanes with a median half-life of approximately 7.1 y (95% quantile: (2.8, 8.8)) for H1N1pdm09 and 3.5 y (95% quantile: (1.4, 5.2)) for H3N2 (Fig. 2 C,D); thus, the duration of protection in children is similar to adults’ against H1N1pdm09, but shorter against H3N2. We find that unlike in adults, infection with H1N1pdm09 generates a long-term boost in titer that is 30% the size of the acute boost (*ζ*_children,*s*_ = 0.3 (95% CI: 0.2, 0.5), Table 1), allowing children to gain long-term protection as their baseline titer eventually rises above the *TP50*_children_ through repeated exposures. In H3N2, by contrast, we estimate only a small long-term boost (*ζ*_children,*s*_ = 0.04 (95% CI: 0.02, 0.07), Table 1), which could reflect the antigenic evolution of circulating strains and the change in the strain used in the HI assay during the study.

### The models reproduce population-level patterns of infection and other estimates of protection

Despite being fitted to individuals’ titers, the models recover reasonable population-level patterns of infection for both subtypes. From the simulated latent infections, we inferred the incidence in children and adults (Fig. 3, Table S2). Because the models assume that the community-level, subtype-specific influenza intensity affects an individual’s infection risk (Methods, Eq. 7), it is unsurprising that periods of high incidence in the simulated study population match those in the community (Fig. 3). However, the absolute incidence in the study population is effectively unconstrained, emerging from the estimated subtype-specific scaled transmission rate, *β*_scaled,*s*_, and protection parameters. The results nonetheless match estimates from other populations. The incidences of individual H1N1pdm09 epidemics range from 7-12% in adults and 10-17% in children (Table S2). For H3N2, the incidences range from 4-12% in adults and 6-23% in children. Estimates of seasonal influenza incidence in the United States are 5-20% based on combined serology and viral infection (of influenza A and B) [27] and 3-11% based on symptomatic PCR-confirmed infections of influenza A [28]. Converted to annual rates, the simulated incidences are similar, ranging from 5-17% in adults and 7-29% in children.

**Figure 3:**
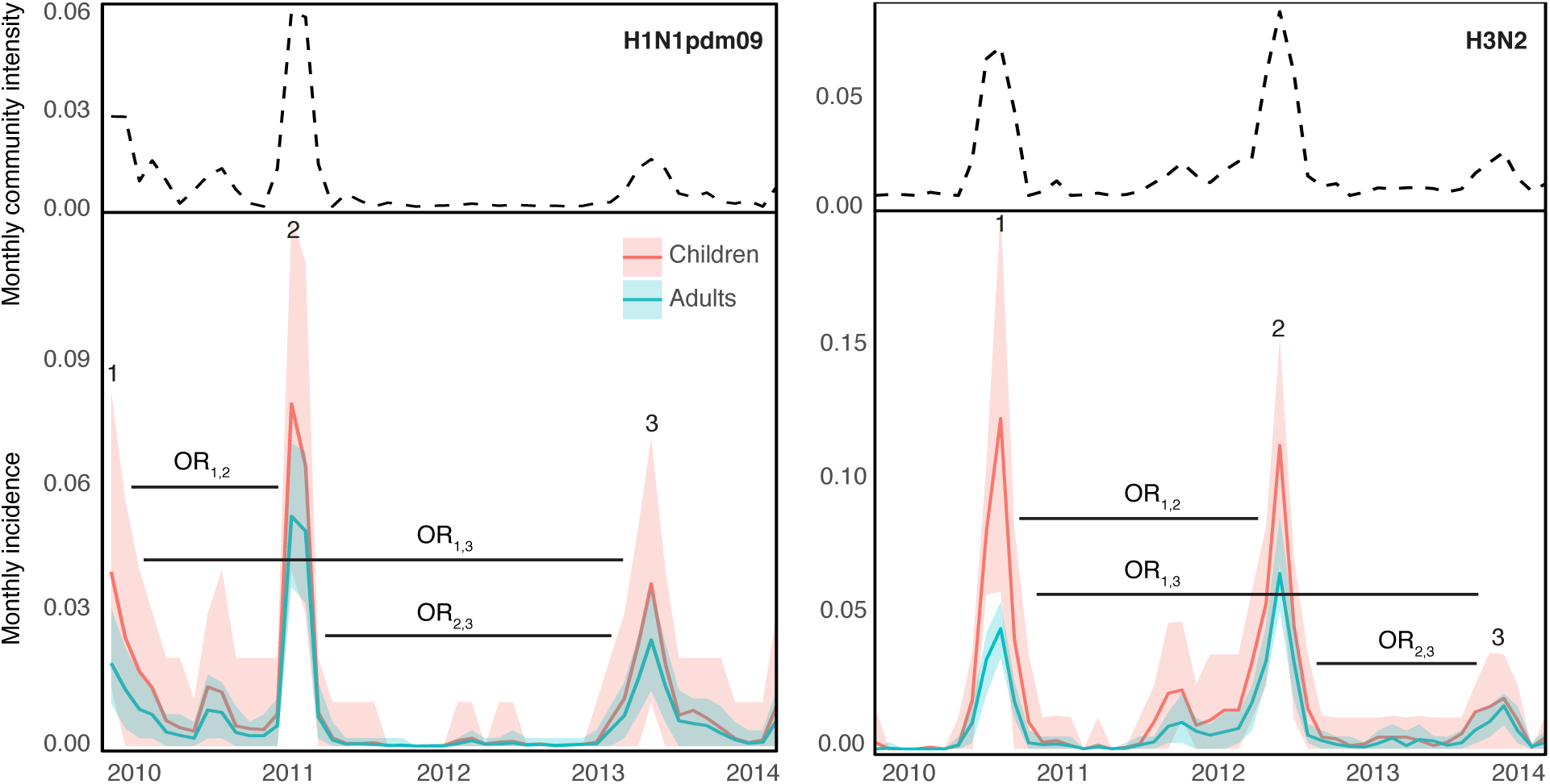
Simulated monthly incidence for H1N1pdm09 (**a**, bottom) and H3N2 (**b**, bottom) in children and adults, averaged over 1000 simulations, contrasted with respective monthly community intensities (**a** and **b**, top). The shaded areas are bounded by the 2.5% and 97.5% quantiles from the simulations. Horizontal black bars denote inter-epidemic periods for odds ratios (OR).

We estimate that for both subtypes, the daily within-household intensity (the increased risk of transmission given that at least one member in the household is infected) is roughly 5%, or ~ 25% over an average five-day infectious period (*ω*_*s*_, Table 1). This is consistent with published estimates of secondary household attack rates for influenza A virus ranging from 10% to 30% [29, 30, 31, 32, 33, 34]. For example, for a young child at the peak of a H1N1pdm09 epidemic, the daily risk of transmission from the community is approximately 2.1% per day. Therefore, our results suggest that the within-household transmission rate is at least twice as high as the maximum community-level risk. The same is true for H3N2.

The simulated infections reproduce other estimates of protection over time. We estimated the odds ratios of protection between epidemics (Table 2). We find evidence of inter-epidemic H1N1pdm09 protection for children between 2009 and 2011, consistent with a previous analysis of this trial that used ≥ 4-fold titer rises to indicate infection [19], and between 2011 and 2013. We also find evidence of protection for adults for the same two inter-epidemic periods. Protection against H3N2 in both children and adults occurred between 2010 and 2012 and between 2012 and 2013. The point estimates of the odds ratios inferred from 4-fold titer rises in children are lower than those inferred from latent infections in this model, suggesting that past infection may protect more against large titer rises than infection per se.

**Table 2:** Inter-epidemic odds ratios of infection, predicted from 1000 replicate simulations of the models for H1N1pdm09 and H3N2 at the MLEs. Epidemics are numbered as in Fig. 3. The rightmost column shows odds ratios estimated using *≥* 4-fold changes in titer as an indicator of infection.

| Subtype |  |  | OR [95% quantiles] | Estimate from [19] |
| --- | --- | --- | --- | --- |
| H1N1pdm09 | Adults | OR<sub>1,2</sub> | 0.55 [0.51, 0.70] |  |
|  |  | OR<sub>2,3</sub> | 0.72 [0.69, 0.80] |  |
|  |  | OR<sub>1,3</sub> | 0.86 [0.74, 1.16] |  |
|  | Children | OR<sub>1,2</sub> | 0.39 [0.12, 0.85] | 0.27 [0.10, 0.76] |
|  |  | OR<sub>2,3</sub> | 0.29 [0.08, 0.65] |  |
|  |  | OR<sub>1,3</sub> | 0.48 [0.09, 1.10] |  |
| H3N2 | Adults | OR<sub>1,2</sub> | 0.37 [0.19, 0.56] |  |
|  |  | OR<sub>2,3</sub> | 0.55 [0.38, 0.76] |  |
|  |  | OR<sub>1,3</sub> | 0.81 [0.53, 1.03] |  |
|  | Children | OR<sub>1,2</sub> | 0.48 [0.10, 0.89] | 0.39 [0.18, 0.83] |
|  |  | OR<sub>2,3</sub> | 0.28 [0.06, 0.79] |  |
|  |  | OR<sub>1,3</sub> | 0.85 [0.82, 1.09] |  |

### No effect of group-level HA imprinting or heterosubtypic infection on susceptibility

Previous work has suggested that primary infection with a subtype reduces susceptibility to severe disease and death with related subtypes, a phenomenon known as imprinting [8, 35]. Influenza A HAs fall into two phylogenetic groups, with H1 and H2 belonging to Group 1 and H3 to Group 2. We estimated the protective effect *α*_imp,*s*_ of primary infection with a subtype of one HA group on the rate of infection with subtype *s* of the same HA group by

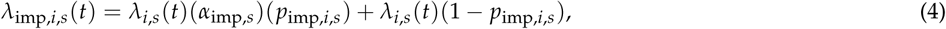

where *λ*_imp,*i*,*s*_ (*t*) is the force of infection on individual *i* with subtype *s* at time *t* considering imprinting, and *λ*_*i*,*s*_ (*t*) is the baseline infection rate for an individual not imprinted with that subtype’s HA group (Eq. 6). The probability of having had a primary infection with the same group as subtype *s* is *p*_imp,*i*,*s*_, and *α*_imp,*s*_ is thus the change in that baseline force of infection from imprinting. Imprinting implies *α*_imp,*s*_ < 1. We calculate *p*_imp,*i*,*s*_ based on the individual’s birth date, the current date, and historical incidence data (Methods, Fig. S3A). Birth-year effects and age-specific effects are often confounded, but are potentially distinguishable in longitudinal data from individuals of similar ages but different primary exposures. We therefore fit the imprinting models for H1N1pdm09 and H3N2 to data from middle-aged adults (35-50 y), whose first exposures were to Group 1 (mainly H2N2) or Group 2 (H3N2) viruses (Fig. S3B). For H3N2, we thus estimate the effect of homosubtypic imprinting, and for H1N1pdm09, we estimate group-level imprinting from primary infection with either H1N1 or H2N2 (Table S3). The 95% confidence intervals for the maximum likelihood estimates of the imprinting effect (Fig. S3C) show the model is consistent with Group 1 protection ranging from 0.6 to 1.0, i.e., 0-40% reduction in susceptibility, and with Group 2 protection between 0.9 and 1.2, suggesting no protection. Epidemiological and immunological studies have suggested that infection with one subtype might protect in the short term against another [36, 37, 38]. To estimate the duration of heterosubtypic protection, we fitted a two-subtype model of H1N1pdm09 and H3N2, fixing the parameters that govern homosubtypic immunity at the MLEs of the best-fit single-subtype models (Table 1). Let *q*_homosubtypic,*i*,*s*_ denote susceptibility to subtype *s* determined only by homosubtypic protection. Heterosubtypic protection after infection with subtype *m /*= *s* contributes to the susceptibility against subtype *s* such that the net susceptibility to subtype *s*, *q*_*i*,*s*_ (*t*), is

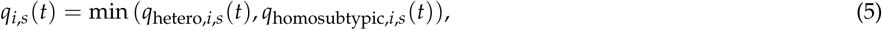

where *q*_hetero,*i*,*s*_ (*t*) is determined by the time since infection with subtype *m* (Eq. 17). We assumed the rate of waning of heterosubtypic protection, *w*_nonspecific,*m*_, is identical for both subtypes. In these data, the model estimates that any heterosubtypic protection is fleeting (Fig. S4; half-life from *w*_nonspecific,*m*_ = 0.002 y; 95% CI: 0.0, 0.07).

### Model validation and sensitivity analysis

In addition to comparing the models’ results to other estimates of population-level incidence and protection between epidemics, we investigated the model’s ability to match other features of the data. The best-fit models reproduce the observed distributions of 1-, 2-, and 4-fold titer rises, considering all individuals’ trajectories together (section S2.1; Figs. S7 and S6). However, the models tend to overestimate how much an individual’s titer varies over time (section S2.2; Figs. S8 and S9). This suggests the model might not be fully capturing individual heterogeneity in infection risk and/or the response to infection, although individuals’ trajectories appear reasonable by eye (Figs. S10, S11). Beyond estimating the factors that correlate with protection, we examined the robustness of our model to other assumptions. Our results are robust to changes in the initial conditions, namely, how recently individuals are assumed to have been infected (section S2.3). Results also do not change with an alternate scaling of the community influenza intensity to account for increased surveillance during the 2009 H1N1pdm09 pandemic (section S2.4). Additionally, our assumptions about the measurement error are consistent not only with values estimated by others [17, 39] but also with the error estimated from replicate titer measurements in the data (section S2.5).

### Discussion

Our results suggest that protection against influenza A has different origins in adults and children. In children, the HI titer is a good correlate of protection, and infection durably boosts titers against H1N1pdm09. In adults, time since infection is the better correlate of protection, and infection is associated with small to no long-term changes in titer. These results suggest that children tend to produce antibodies that target the head of the HA, which could mediate protection, whereas adults rely on antibody responses to other sites or potentially other forms of immunity for protection. This model is consistent with the concepts of antigenic seniority and original antigenic sin. Antigenic seniority refers to the phenomenon that individuals’ highest antibody titers to influenza are to strains that circulated in childhood [22], and original antigenic sin is the process by which antibody responses to familiar sites are preferentially reactivated on exposure to new strains [5, 7, 6]. With time, these familiar sites may be the ones that are most conserved. On HA, these sites would tend to be away from the fast-evolving epitopes near the receptor binding domain, and would not be readily detected by HI assays. Consistent with this view, epidemiological studies have shown that levels of stalk-directed antibodies increase with age [40, 41]. Our model shows that these differences are persistent and relate to protection.

We estimated that protection in both children and adults wanes with an average half-life of 3.5-7 years, lasts longer against H1N1pdm09 than H3N2, and lasts slightly longer in adults compared to children against H3N2. These timescales are consistent with the estimated decay of immunity over 2-10 years due to antigenic evolution in population-level models [42, 43]. In contrast to adults, the dependence of protection on HI titer in children leads to substantial variation in susceptibility over time (Fig. 2). This heterogeneity may well extend to adults but is difficult to identify without much longer time series or other immune assays. The models’ tendency to overestimate individuals’ titer variation over time suggests that important differences between individual responses could be missing (Figs. S8 and S9). Again, longer observation periods and more complete observations of the immune response can help separate these factors from behavioral or environmental differences in infection risk.

We find minimal evidence that HA imprinting and heterosubtypic immunity affect susceptibility to infection. Previous analyses of HA group-level imprinting have suggested that imprinting reduces the rate of severe disease and death [8]. Serological testing in this study occurred independent of symptoms. If the model were estimating protection from symptomatic, medically attended infections instead, a stronger effect might have been supported. In the same vein, heterosubtypic immunity, for which there is good evidence [36], might reduce the severity of illness rather than prevent infection [44, 45]. Another possibility is that the discordance of H1N1pdm09 and H3N2 epidemic peaks in this study (Figs. 3 and S12) reduced the model’s power to detect short-term cross-protection (Fig. S4).

This work has several limitations. We return again to the concept of heterogeneity. Though our models support substantial variability in the short-term titer boost after infection, our data lack multiple PCR-confirmed infections from the same people. Thus, we cannot distinguish the nonspecific variability at each infection (*σ*_*ai*,*s*_) from consistent differences between individuals, which might be expected if people persistently target different sites on HA, NA, and other proteins. However, the statistically indistinguishable acute titer boosts in children and adults indicate that despite adults’ low baseline HI titers, we were not systematically underestimating their rate of infection. Although our results provide insight into differences between children and adults, we have obviously not modeled the evolving response in individuals over a lifetime, spanning infancy to old age. We thus cannot resolve how age-related phenomena interact with immune history to affect the response to infection.

Broadly, our results estimate several years of protection after infection with influenza A in children and adults, and suggest that this protection is associated with different immune responses, which are consistent original antigenic sin. These results underscore the need for a deeper understanding of the factors that determine the variable response to infection among individuals, and for better correlates of immune protection. They also underscore the utility of longitudinal cohorts and mechanistic models to investigate the dynamics of influenza.

## Methods

### Study description

The data are part of a community-based study of influenza virus infections in households that was conducted in Hong Kong between 2009 and 2014 [19]. The study tracked individuals in 796 households, each of which included at least 1 child aged 6-17 y that had no contraindications against the trivalent inactivated influenza vaccine (TIV). One eligible child 6-17 y of age per household was randomized to receive either a single dose of TIV or saline placebo, regardless of influenza vaccination history. In vaccinated individuals, sera was collected at baseline prior to vaccination (August 2009 - February 2010) and 1 month after vaccination. In all individuals, sera was collected after enrollment in the autumn of 2009 and again each subsequent autumn, and each spring for at least 25% of participants. Participants were invited annually to continue enrollment. Individuals reported receipt of the influenza vaccine outside of the trial annually.

Participants and household contacts were encouraged to record systemic and respiratory symptoms daily in diaries. Acute respiratory infections (ARIs) were surveilled by telephone calls every 2 weeks, and households were encouraged to report ARIs promptly to the study hotline. Home visits were triggered by the presence of any 2 the following: fever (37.8*◦*C), chills, headache, sore throat, cough, presence of phlegm, coryza, or myalgia in any household member. Combined nasal and throat swabs were collected from all household members during home visits, regardless of illness.

### Ethical approval

The study protocol was approved by the Institutional Review Board of the University of Hong Kong. All adults provided written consent. Parents or legal guardians provided proxy written consent for participants ≤17 y old, with additional written assent from those 8–17 y.

### Laboratory testing

Serum specimens were tested by HI assays in serial doubling dilutions from an initial dilution of 1:10 [19, 18]. The antibody titer was taken as the reciprocal of the greatest dilution that gave a positive result. Sera from year 1 (2009–2010) and year 2 (2010–2011) were tested against A/California/7/2009(H1N1) and A/Perth/16/2009-like (H3N2). In years 3-5 (2011–2012, 2012–2013, and 2013–2014), sera were tested against the same H1N1pdm09 strain and against A/Victoria/361/2011-like (H3N2). Sera from consecutive years were tested in parallel, such that duplicate titer measurements exist for sera sampled during the middle of the study. For this analysis, we used the first titer measurement obtained for any serum sample. Nose and throat swabs were tested by reverse transcription polymerase chain reaction (PCR) for influenza A and B viruses using standard methods, as described previously [46].

### Data included in this analysis

We fitted models to HI titers from 706 individuals (including 114 children ≤15 y old at enrollment) from 337 different households that were not vaccinated as part of the study and reported no vaccination at any season during follow-up. We excluded individuals with any missing vaccination information. Individuals in this subset were sampled at a median of 6.6 months over a median 5.0 years of follow-up. Households ranged in size from 1 to 5 members (median = 2 members). Figure 4,c gives the number of samples over the study among children and adults. Among children, the median age at enrollment was 11 y, and the age range was 3–15 y. Among adults, the median age of enrollment was 43 y, the age range was 16–77 y, and 89% of adults were between 25 and 55 y. We fitted sub-models to data from 50 individuals (including 29 children ≤15 y old at enrollment) with PCR-confirmed H3N2 infection and 78 individuals (including 42 children ≤15 y old at enrollment) with PCR-confirmed H1N1pdm09 infection (section 1.1). No individuals had multiple PCR-confirmed infections. The data for this analysis are the date of the subtype-specific PCR-positive nasal swab and the closest titer measurements surrounding the positive swab. For H3N2, the median time between the pre-infection titer measurement and the PCR-positive swab was 5.3 months, and the median time between the PCR-positive swab and the post-infection titer measurement was 2.6 months. For H1N1pdm09, the median time between the pre-infection titer measurement and the PCR-positive swab was 2.4 months, and the median time between the PCR-positive swab and the post-infection titer measurement was 6.6 months. Figure 4d shows the number of PCR-confirmed infections by subtype in adults and children over time.

**Figure 4:**
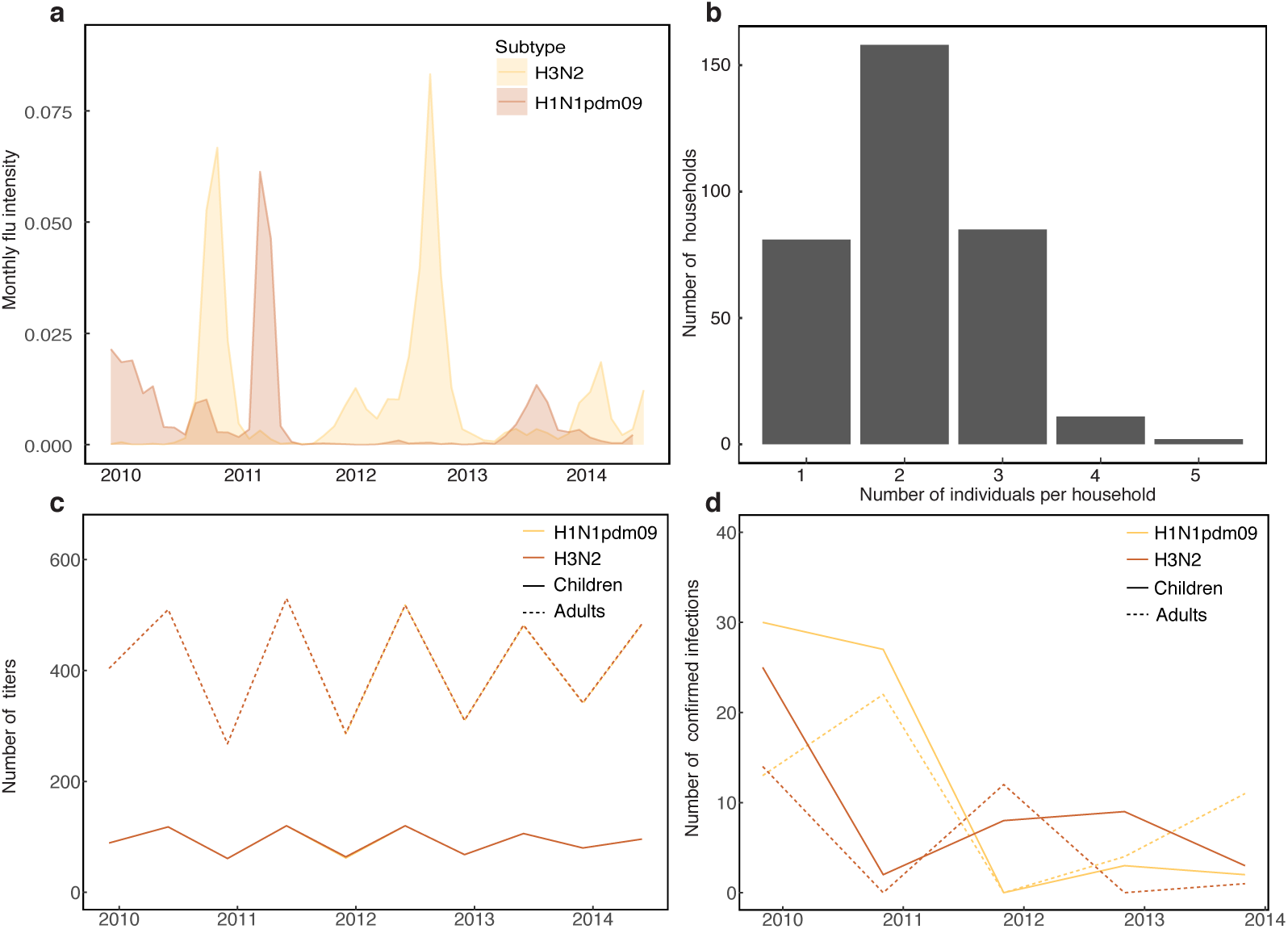
Features of the data. **(a)** Community intensity of H3N2 and H1N1pdm09 (ILI x % influenza-positive) over the study. **(b)** Distribution of household sizes in the study cohort. **(c)** Number of titer samples for H3N2 and H1N1pdm09 in children and adults over the study. **(d)** Number of PCR-confirmed infections with H3N2 and H1N1pdm09 in children and adults over the study.

### Complete model description

The model simulates the titer and infection dynamics for each individual. Briefly, an individual’s instantaneous infection rate or force of infection, *λ*_*i*,*s*_ (*t*), is determined by the rate of exposure to infectious contacts in the community and household and by the individual’s susceptibility, defined as the probability of infection per infectious contact. Infections occur stochastically and can change the latent titers. Simulating the model generates a latent titer for each individual at each observation. The measurement model then calculates the likelihood of each observed titer given the contemporaneous latent titer. The log likelihood of the model under any set of parameters is then the sum of the log likelihoods across individuals, which are the sums of the log likelihoods across observations. The titer and infection dynamics are described in this section.

#### 1. Exposure to infection

The instantaneous per capita infection rate for individual *i* with subtype *s* is denoted *λ*_*i*,*s*_ (*t*). An individual’s overall rate of infection with subtype *s* comprises a rate of infection from the community, *λ*_community,*i*,*s*_, and another from the household, *λ*_household,*i*,*s*_,

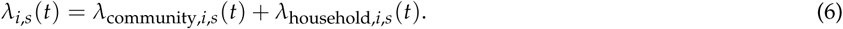

The rate of exposure to infection in the community is influenced by age-specific contact rates [20, 12], the age distribution of infectious contacts [47], and a proxy for the prevalence of the subtype in the community. The community-driven rate of infection is the rate of exposure modified by the individual’s susceptibility to that subtype,

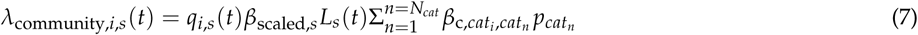

where *q*_*i*,*s*_ (*t*) is the individual’s susceptibility to that subtype (or per-infectious-contact probability of infection), *β*_c,*cati*, *catn*_ is the fixed contact rate for an individual of age category *cat*_*i*_ with individuals of age category *cat*_*n*_ (Table S4, section S2.6), *N*_*cat*_ is the number of contact age categories (5 in total, Table S4), *L*_*s*_ (*t*) is a proxy of influenza activity for subtype *s*, and *p*_*catn*_ is the fraction of influenza infections attributable to age class *cat*_*n*_. The parameter *β*_scaled,*s*_ scales the flu intensity to determine the per-infectious-contact transmission rate at time *t*. We calculate *L*_*s*_ (*t*) from weekly community surveillance data as (ILI/total general practitioner consultations)(% specimens positive for subtype *s*) (Fig. 4) [48]. We impose a minimum threshold min(*L*_*s*_ (*t*)) = 10^*−*5^.

Individual *i* also experiences infection risk from household members,

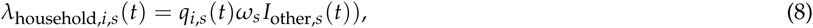

where *q*_*i*,*s*_ (*t*) is the individual’s susceptibility, and *I*_other,*s*_ (*t*) is an indicator variable that equals 1 if any other household member is infected with subtype *s* at time *t* and 0 if not. Because we do not track households in their entirety (we sample only individuals that were not vaccinated during the study), we do not model density-dependent within-household transmission. Rather, *ω*_*s*_ describes the daily influenza exposure rate that individual *i* experiences with the presence of any infected family member.

#### 2. Susceptibility to infection based on HI titer to the infecting strain, non-HI-correlated protection, or both

Individual *i*’s susceptibility to subtype *s* at time *t*, *q*_*i*,*s*_ (*t*), is defined as the probability of infection given contact with an infected. Complete protection corresponds to *q*_*i*,*s*_ (*t*) = 0, complete susceptibility to *q*_*i*,*s*_ (*t*) = 1, and 0 < *q*_*i*,*s*_ (*t*) < 1 to partial protection. We use two base functions to model susceptibility. One function assumes susceptibility depends on the HI titer against the infecting strain (the HI-correlated component), and the other on the time since infection with that subtype (the non-HI-correlated-component). The HI-correlated component of susceptibility, *q*_1*i*,*s*_ (*t*), is a logistic function of the HI titer [11, 49] (Fig. 1, Step 1b). Because previous studies suggest that the relationship between titer and susceptibility changes with age [13], we estimate the relationship separately for children and adults. The HI-correlated susceptibility of individual *i* to subtype *s* at time *t*, *q*_1*i*,*s*_ (*t*), is given by the logistic function,

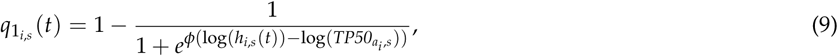

where *h*_*i*,*s*_ (*t*) is the latent titer and *TP50_ai_*_,*s*_ is the subtype- and age-specific 50% protective titer. The scaling parameter *φ*, which determines the shape of the logistic curve, is fixed (Table S4). The non-HI-correlated component of susceptibility, *q*_2*i*,*s*_ (*t*), assumes initially complete protection that wanes at a constant rate after infection,

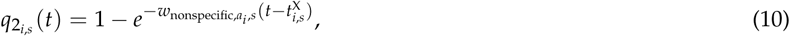

where *w*_nonspecific,*ai*, *s*_ is the rate of waning, fitted separately for children and adults, and 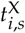 is the time of infection.

The susceptibility *q*_*i*,*s*_ (*t*) is the weighted average of the two components (Eq. 3).

#### 3. Titer dynamics after infection

Infections can affect the latent titers. We model the post-infection titer dynamics as a series of equations that describe the acute boost, the waning from peak titer, and the potential long-term titer boost. Together, these equations determine the latent titer of individual *i* against subtype *s* at any time *t*, *h*_*i*,*s*_ (*t*).

When individual *i* is infected with subtype *s*, antibody titers increase from the time of infection and eventually peak. The acute boost occurs occurs according to *f*_rise_,

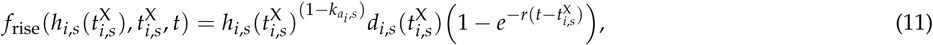

where 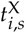 and *hi*,*s* (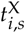) give the time and titer, respectively, of the most recent infection; *r* gives the fixed rate of titer rise after infection (Table S4); and *d*_*i*,*s*_ (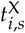) is the magnitude of the short-term boost. Recall that the time 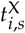 of a simulated infection is driven by the individual’s infection rate, *λ*_*i*,*s*_ (*t*). The age- and subtype-specific parameter *k*_*ai*,*s*_ determines the dependence of the titer boost on the pre-infection titer. When positive, it allows for an antibody ceiling effect [50], whereby higher pre-infection titers have smaller boosts (section S1.1).

Multiple studies demonstrate heterogeneity in the short-term titer rise after infection [51, 39]. Therefore, we allow for variability in the magnitude of the short-term boost for each infection,

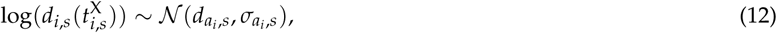

where *d*_*ai*,*s*_ and *σ*_*ai*,*s*_ give the age- and subtype-specific log mean and standard deviation, respectively, of the boost. We estimate the parameters *k*_*ai*,*s*_, *d*_*ai*,*s*_, and *σ*_*ai*,*s*_ from a sub-model fitted to data from individuals with PCR-confirmed infection (section S1.1). We then fix the values of these parameters in the main model.

After peaking at time 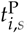, the titer wanes exponentially at a fixed rate *w* (Table S4) to an individual’s subtype-specific baseline titer, *h*_baseline,*i*,*s*_ (*t*). Therefore, the titer after the peak short-term response is given by

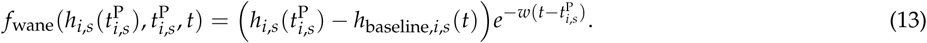

Infection may cause a long-term boost, *d*_longterm,*i*,*s*_, (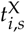), that does not wane, where *d*_longterm,*i*,*s*_ (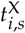) is defined as a fraction *ζ*_*ai*,*s*_ of the acute boost,

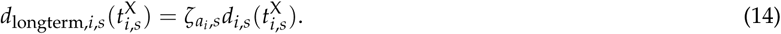

The long-term boost changes the baseline titer after each infection at time 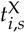,

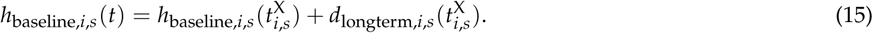

Let *T*_peak_ denote the fixed length of time between infection and peak titer (Table S4). The complete expression for *h*_*i*,*s*_ (*t*) is then

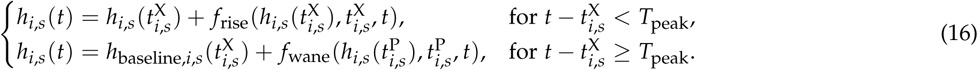

#### 4. Heterosubtypic immunity

Heterosubtypic immunity acts as a non-specific form of protection against subtype *s* following infection with subtype *m* at time 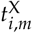 and wanes at rate *w*_nonspecific,*m*_,

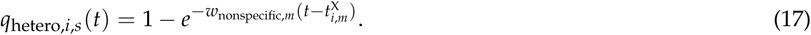

### Measurement model and likelihood function

The observed titer *H*_obs,*i*,*s*_ (*t*) relates to the corresponding latent titer *h*_*i*,*s*_ (*t*) via the measurement model, which accounts for error in the titer measurements and the effect of discretization of titer data into fold-dilutions. The observed titers are fold-dilutions in the range [<1:10, 1:10, 1:20 …, 1:5120]. Consistent with other models [16, 12, 14], we define a log titer (*logH*) for any titer *h*,

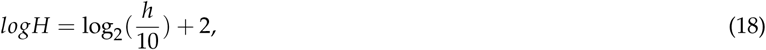

such that the log of the observed titer takes on discrete values in the range [1,11]. In order to relate the observed titer, *H*_obs,*i*,*s*_ (*t*), to the latent titer, *h*_*i*,*s*_ (*t*), we begin by transforming both into log titers (Eq. 18), yielding the log observed titer *logH*_obs,*i*,*s*_ (*t*) and the log latent titer *logH*_*i*,*s*_ (*t*) We assume that the observed log titer to subtype *s* is normally distributed around the log latent titer,

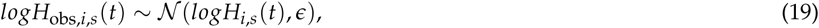

where *ε* gives the standard deviation of the measurement error.

Following other analyses that quantified the measurement error associated with different titers [15, 39], we assign a lower measurement error (*c* = 0.74 log titer units, Table S4) for undetectable (< 10) titers. The observed titer is censored at integer cutoffs, such that the likelihood of observing *logH*_obs,*i*,*s*_ (*t*) = *j* given log latent titer *logH*_*i*,*s*_ (*t*) is

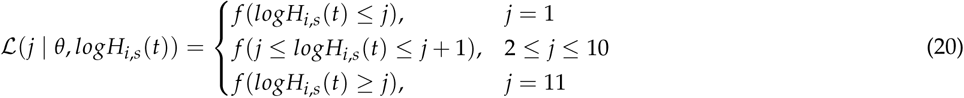

where *θ* gives the parameter vector and *f* is specified as in Eq. 19. For each individual, the log likelihood is the sum of the log likelihoods across observations,

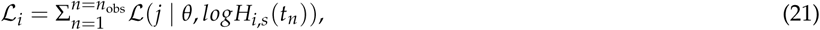

where *n*_obs_ is the number of observations for individual *i* and *t*_*n*_ gives the time at observation *n*. The log likelihood of the model for any parameter set *θ* is then the sum across *n*_ind_ individuals,

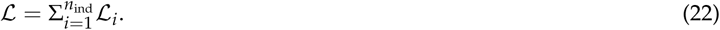

Table S4 summarizes the model parameters and state variables.

### Initial conditions

We assign each individual’s initial latent subtype-specific baseline titer, *h*_baseline,*i*,*s*_ (0), based on that individual’s lowest observed titer during the study, 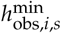. Because an observed HI titer represents the lower bound of a two-fold dilution, we draw *h*_baseline,*i*,*s*_ (0) for each realization of the model according to

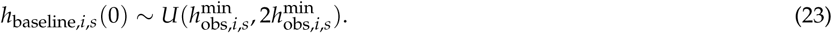

The values of the initial latent titer, *h*_*i*,*s*_ (0), and the initial susceptibility, *q*_*i*,*s*_ (0), depend on the time of most recent infection, which may have occurred before entry in the study. To initialize the latent states for each individual, we draw the time of the most recent infection from the density of subtype-specific flu intensity, *L*_*s*_ (*t*), in the seven years before the first observation. In this way, we account for known epidemic activity in Hong Kong before the beginning of the study (Fig. S5). For children less than 7 y old, the distribution is truncated at birth, and the density includes the probability that the child is naive to influenza infection. For sensitivity analysis, we fitted the models using other assumptions about the density from which we drew the time of most recent infection (section S2.3 and S2.4).

### Likelihood-based inference

The titer and infection dynamics of each individual are modeled as a partially observed Markov process (POMP). The model for each subtype is a “panel POMP” object, or a collection of the individual POMPs with shared age- and subtype-specific parameters. We use panel iterated filtering (PIF) to fit the models [52, 53]. Iterated filtering uses sequential Monte Carlo (SMC) to estimate the likelihood of an observed time series. In SMC, a population of particles is drawn from the parameters of a given model to generate Monte Carlo samples of the latent dynamic variables. To evaluate the likelihood of a parameter set, SMC is carried out over the time series for each individual, generating a log likelihood for the corresponding panel unit. The log likelihood of the panel POMP object is the sum of the individuals’ log likelihoods. Within each PIF iteration, filtering, or weighted particle re-sampling, occurs once for all observations from each individual. One PIF iteration is one pass of the weighted re-sampling over all individuals. Damped perturbations to the parameters occur between iterations. As the amplitude of the perturbations decreases, the algorithm converges to the maximum likelihood estimate [52]. For each model, we initialize the iterated filtering with 100 random parameter combinations. We perform series of successive 50-iteration MIF searches, with the output of each search serving as the initial conditions for the subsequent search. We use 10,000 particles for each MIF search. The likelihood of the output for each search is calculated by averaging the likelihood from ten passes through the particle filter, each using 20,000 particles. We repeat the routine until additional operations fail to arrive at a higher maximum likelihood. For model selection in the sub-model of the acute titer boost, we used the corrected Akaike Information Criterion (AICc) (Table S1) [54]. We obtained maximum likelihood estimates for each parameter and associated 95% confidence intervals by constructing likelihood profiles. We used Monte Carlo Adjusted Profile methods [53] to obtain a smoothed estimate of the profile (section S3).

### Calculating imprinting probabilities

We calculate the probability that an individual’s first influenza A infection was with a particular subtype (H1N1, H3N2, or H2N2) or that the individual was naive to infection at each year of observation. We assume that the first infection occurred between the ages of 6 months and 12 years, as infants are protected by maternal antibodies for the first six months of life [55, 56] Following the original imprinting model by Gostic and colleagues [8], we estimate the probability that an individual with birth year *i* has his or her first influenza A infection in calendar year *j*:

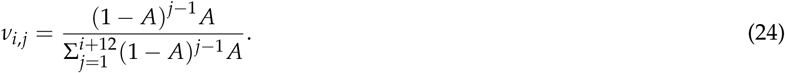

Here, *A* is the constant annual attack rate in seronegative children as estimated by Gostic and colleagues (*A* = 0.28, [8]). Given observation year *y*, the probability that individual *i* was first infected in year *j* is

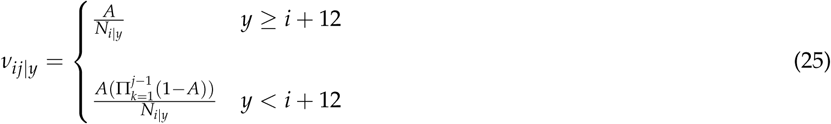

where *N*_*i y*_ is a normalizing factor that enforces the assumption that all individuals have their first infection by age 12 and ensures that all probabilities sum to one for individuals that are ≥12 years old at the observation date. The normalization factor does not apply to individuals < 12 years old, who have some probability of being naive to infection. We combine the probabilities of the age of first infection with annual historical influenza A subtype frequency data (section S2.7) to determine the probability that an individual with birth year *i* had his or her first exposure to subtype *S* in year *j*,

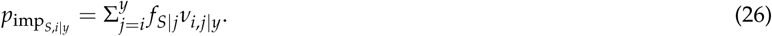

Here, *f*_*S j*_ gives the fraction of specimens of subtype *S* out of all specimens from community surveillance that are positive for influenza A. For individuals younger than 12 years old during the year of observation, the probability that an individual was naive in observation year *y* is

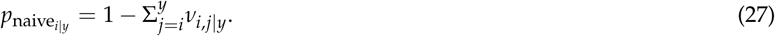

## Code availability

All of the software to run the analysis and produce the figures is available at https://github.com/cobeylab/Influenza-immune-dynamics.

## Data availability

The data for this study are available at https://github.com/cobeylab/Influenza-immune-dynamics.

## Acknowledgements

We are grateful to Phil Arevalo and Ed Baskerville for helpful discussions. We thank the University of Chicago Resource Computing Center for access to high performance computing. This project has been funded in whole or in part with federal funds from the National Institute of Allergy and Infectious Diseases (NIAID), National Institutes of Health (NIH), Department of Health and Human Services, under grant DP2AI117921 (to SC) and CEIRS Contract No. HHSN272201400005C (to SC). The household cohort study was supported by the Research Fund for the Control of Infectious Diseases of the Health, Welfare and Food Bureau of the Hong Kong SAR Government (grant numbers CHP-CE-03, 11100882 and 13120602), the Area of Excellence Scheme of the Hong Kong University Grants Committee (grant number AoE/M-12/06), and the Hong Kong Research Grants Council (grant no. T11-705/14N). Funding was also provided by NIH F30AI124636 to S.R., NIH T32GM007281, and NIGMS grant no. U54GM088558. The funders had no role in study design, data collection and analysis, decision to publish, or preparation of the manuscript.

## Author contributions

S.R., R.S., B.J.C., and S.C. designed the research; S.R. and R.S. performed the analysis; V.J.F., G.M.L., D.K.M., R.A.P.M.P., J.S.M., and B.J.C. provided data and epidemiological expertise; S.R., B.J.C., and S.C. wrote the manuscript. All authors contributed to critical review of the manuscript.

## Competing interests

B.J.C. has received research funding from Sanofi, and honoraria from Sanofi and Roche.

## Materials and correspondence

Sylvia Ranjeva

## Supporting Information (SI)

**Table S1:** Model comparisons for sub-model of short-term boosting. Models “with *k*” include the antibody ceiling effect (Eq. 11, section S1.1), and models “without *k*” do not (*k*_*ai*,*s*_ = 0).

| Subtype | | Model | Log likelihood (SE) | $\Delta\text{AICc}$ |
| --- | --- | --- | --- | --- |
| H1N1pdm09 | Adults | with $k$ | -79.5 (0.06) | 0 |
| | | without $k$ | -81.7 (0.03) | 2.1 |
| | Children | with $k$ | -91.8 (0.01) | 0 |
| | | without $k$ | -95.9 (0.04) | 5.6 |
| H3N2 | Adults | with $k$ | -45.5 (0.04) | 0 |
| | | without $k$ | -48.6 (0.03) | 3.6 |
| | Children | with $k$ | -77.8 (0.07) | 0 |
| | | without $k$ | -85.5 (0.05) | 13.0 |

**Table S2:** Incidence in each epidemic (Fig. 3). The simulated incidence was estimated from the simulated latent infections. The main and bracketed values give the median and 95% quantiles, respectively, from 1000 replicate simulations of the models at the maximum likelihood estimate. Incidence was also estimated from the frequency of *≥* 4-fold titer consecutive titer rises observed in the data.

| Subtype | | Epidemic | Simulated incidence | From observed $\geq 4$-fold changes |
| --- | --- | --- | --- | --- |
| H1N1pdm09 | Adults | 1 | 0.09 [0.06, 0.16] | 0.05 |
|  |  | 2 | 0.12 [0.09, 0.15] | 0.12 |
|  |  | 3 | 0.07 [0.04, 0.09] | 0.06 |
|  | Children | 1 | 0.14 [0.09, 0.33] | 0.10 |
|  |  | 2 | 0.17 [0.12, 0.24] | 0.19 |
|  |  | 3 | 0.10 [0.05, 0.15] | 0.06 |
| H3N2 | Adults | 1 | 0.12 [0.09, 0.15] | 0.16 |
|  |  | 2 | 0.13 [0.11, 0.16] | 0.15 |
|  |  | 3 | 0.04 [0.03, 0.05] | 0.03 |
|  | Children | 1 | 0.22 [0.15, 0.30] | 0.21 |
|  |  | 2 | 0.23 [0.16, 0.30] | 0.24 |
|  |  | 3 | 0.06 [0.02, 0.13] | 0.03 |

**Table S3:** Maximum likelihood estimates of the group-level imprinting effects (*α*_imp,H1N1pdm09_ and *α*_imp,H3N2_) among individuals ages 35-50 y, with 95% confidence intervals (CIs).

| Infecting Subtype | Imprinting Group | Parameter | Population subset | MLE [95% CI] |
| --- | --- | --- | --- | --- |
| H1N1pdm09 | Group 1 | $(\alpha_{\text{imp,H1N1pdm09}})$ | Ages 35 - 50 y | 0.8 [0.6, 1.0] |
| H3N2 | Group 2 | $(\alpha_{\text{imp,H3N2}})$ | Ages 35 - 50 y | 1.0 [0.9, 1.2] |

**Table S4:**
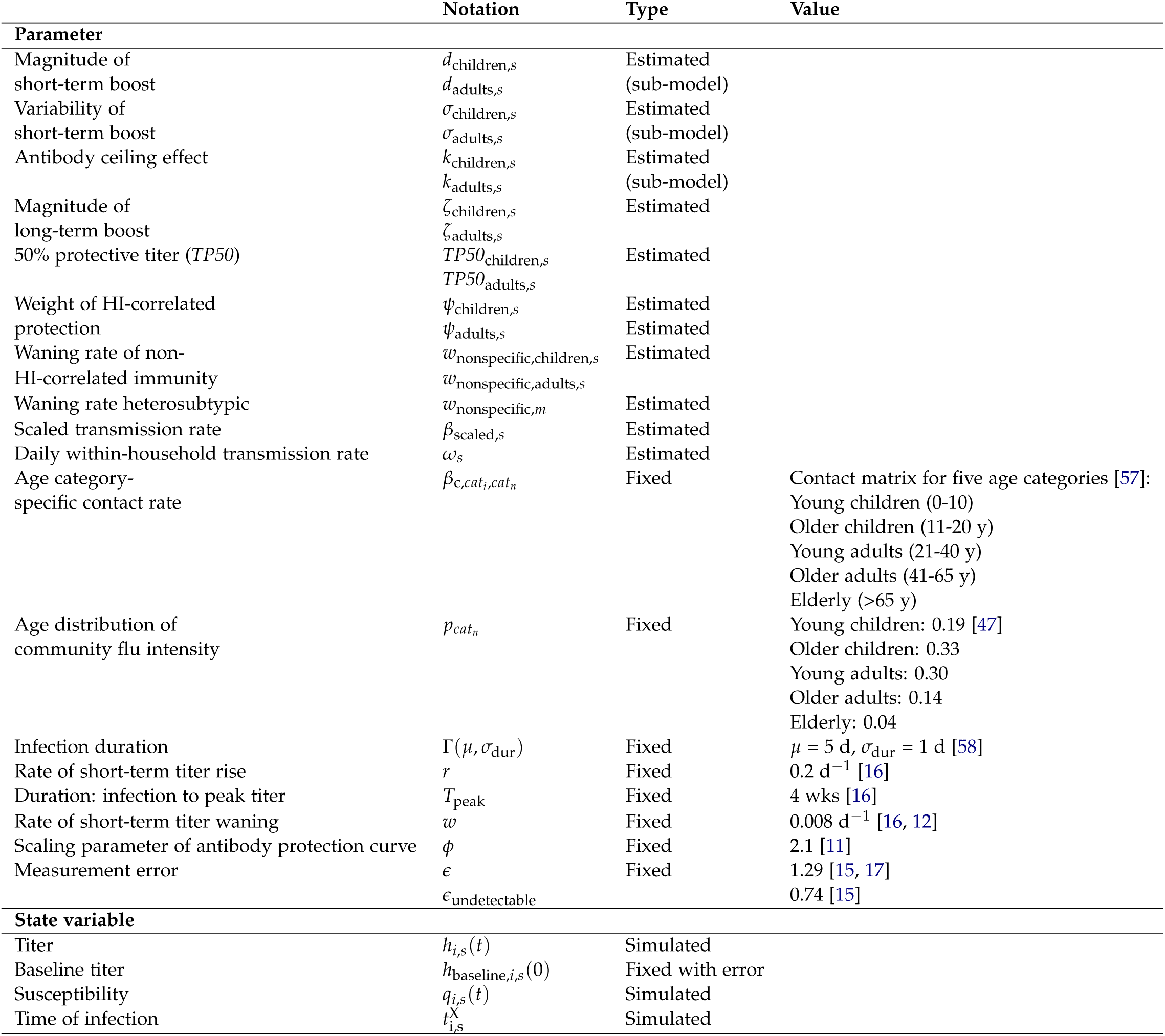
Model parameters and state variables.

**Table S5:** Maximum likelihood estimates of the parameters that govern the short-term titer dynamics with 95% confidence intervals (CIs).

| Subtype |  | Parameter | MLE [95% CI] |
| --- | --- | --- | --- |
| H1N1pdm09 | Adults | $d_{a_i,s}$ | 3.6 [2.4, 5.2] |
| | | $\sigma_{a_i,s}$ | 1.9 [1.4, 2.6] |
| | | $k_{a_i,s}$ | 0.3 [0.1, 1.8] |
| | Children | $d_{a_i,s}$ | 3.5 [3.3, 5.5] |
| | | $\sigma_{a_i,s}$ | 0.9 [0.7, 1.5] |
| | | $k_{a_i,s}$ | 0.6 [0.3, 1.2] |
| H3N2 | Adults | $d_{a_i,s}$ | 4.6 [3.1, 7.1] |
| | | $\sigma_{a_i,s}$ | 1.4 [0.9, 2.3] |
| | | $k_{a_i,s}$ | 1.0 [0.6, 2.5] |
| | Children | $d_{a_i,s}$ | 5.1 [3.9, 7.2] |
| | | $\sigma_{a_i,s}$ | 1.5 [1.0, 2.2] |
| | | $k_{a_i,s}$ | 0.5 [0.3, 1.0] |

**Table S6:** Fraction of children and adults with symptomatic infections (defined by an ARI in the two weeks before PCR-confirmed infection) and primary infections (defined by the absence of infection with or without ARI symptoms in other household members in the two weeks before PCR-confirmed infection) for H1N1pdm09 and H3N2. ARI was defined as having least two of the following symptoms: fever 37.8*◦* C, cough, sore throat, runny nose, headache, myalgia, and phlegm.

| Subtype |  | % Symptomatic infections | % Primary infections |
| --- | --- | --- | --- |
| H1N1pdm09 | Adults | 69.4% | 38.9% |
|  | Children | 64.3% | 54.8% |
| H3N2 | Adults | 76.2% | 38.1% |
|  | Children | 69.0% | 75.8% |

**Table S7:** Maximum likelihood estimate (log scale) of the subtype-specific scaled transmission rate, *β*_scaled,*s*_, for each subtype, with 95% confidence intervals (CIs).

| Subtype | Parameter | MLE [95% CI] |
| --- | --- | --- |
| H1N1pdm09 | $\beta_{\text{scaled},s}$ | -2.8 [-3.0, -2.7] |
| H3N2 | $\beta_{\text{scaled},s}$ | -3.2 [-3.3, -3.1] |

**Figure S1:**
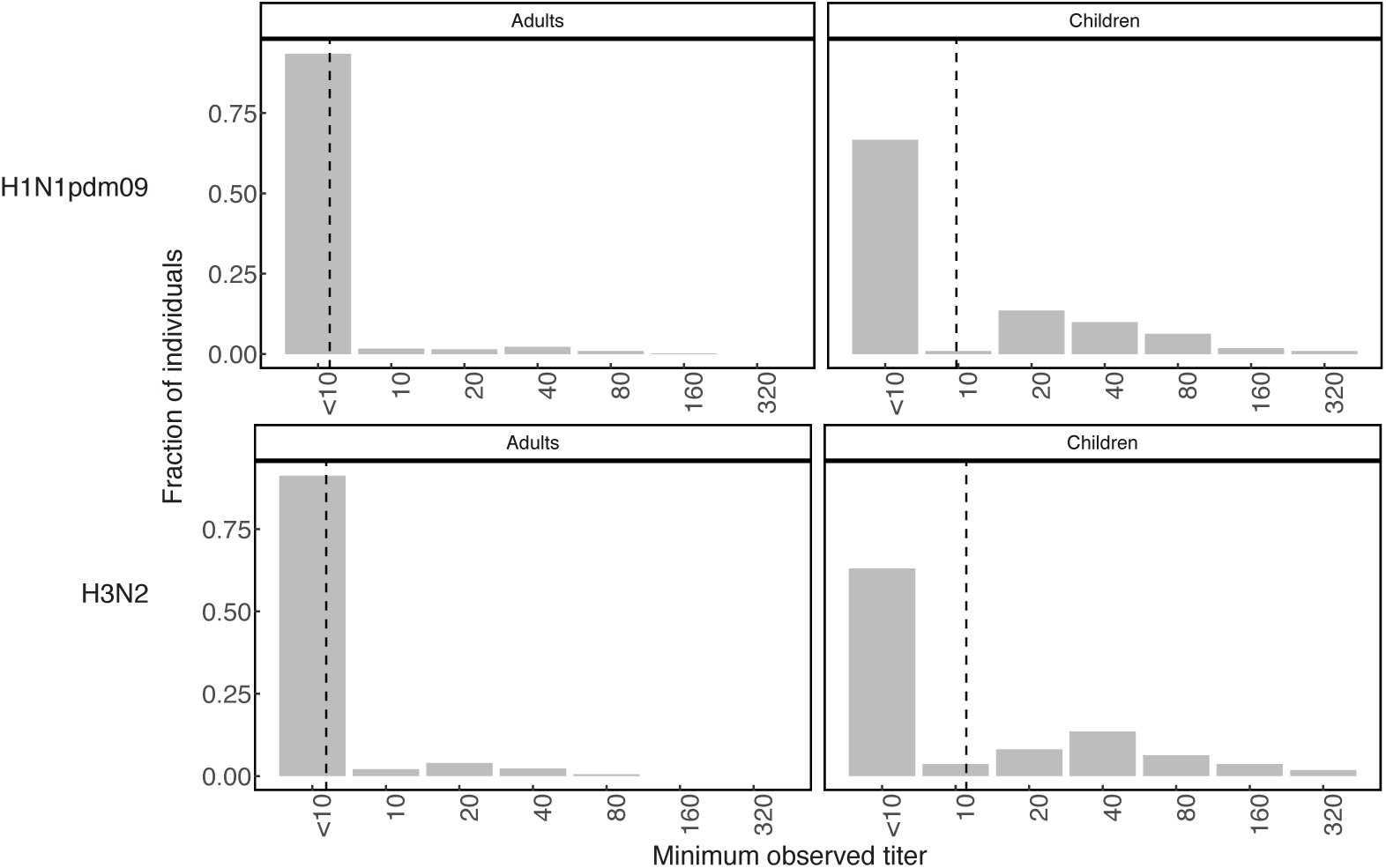
Distribution of the minimum observed (baseline) titers in children and adults for H1N1pdm09 and H3N2. The vertical dashed line gives the geometric mean baseline titer.

**Figure S2:**
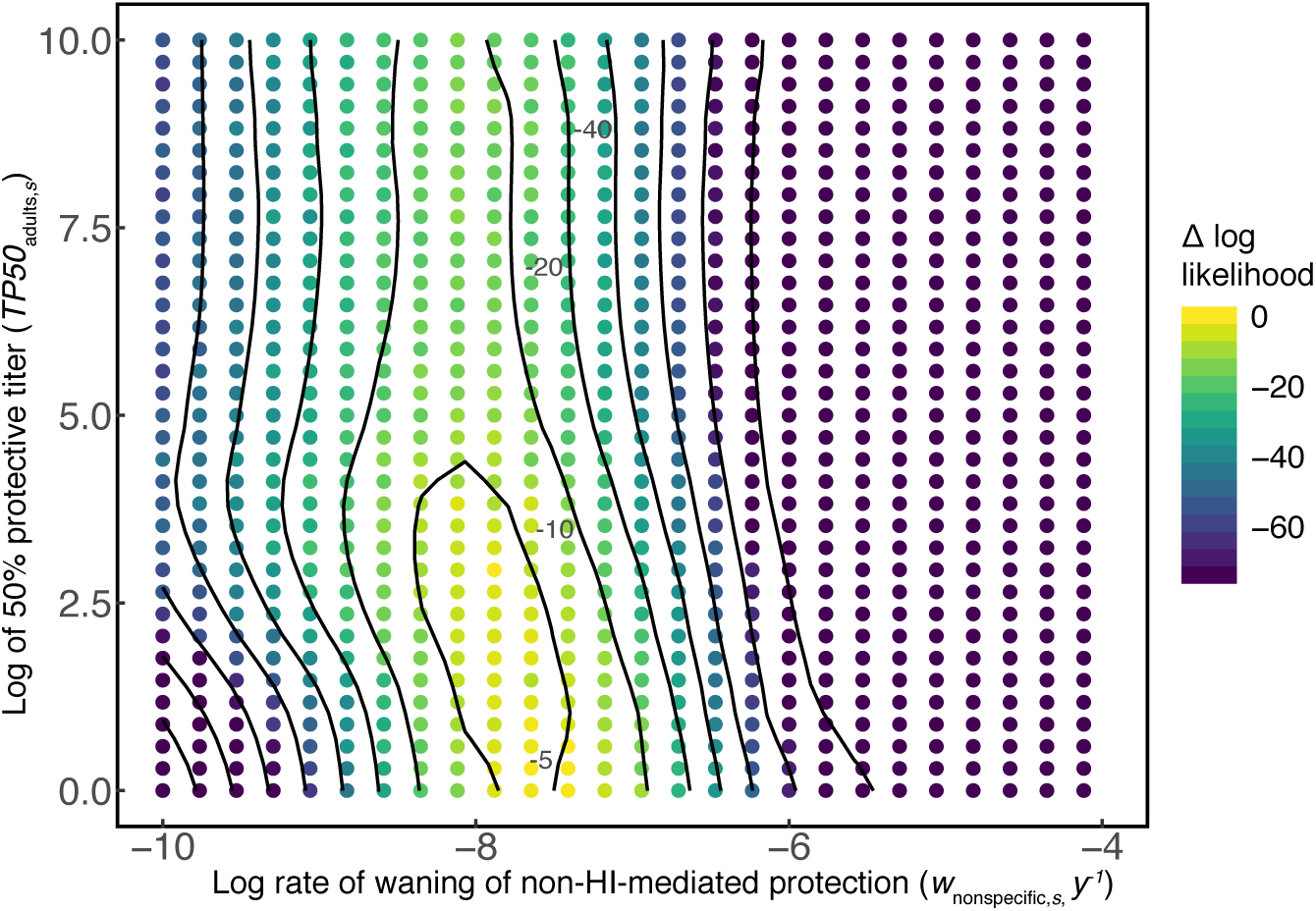
Bivariate likelihood profile of the rate of waning of non-HI-mediated protection in adults and the 50% protective titer in children (*TP*_50,adults_) for H1N1pdm09.

**Figure S3:**
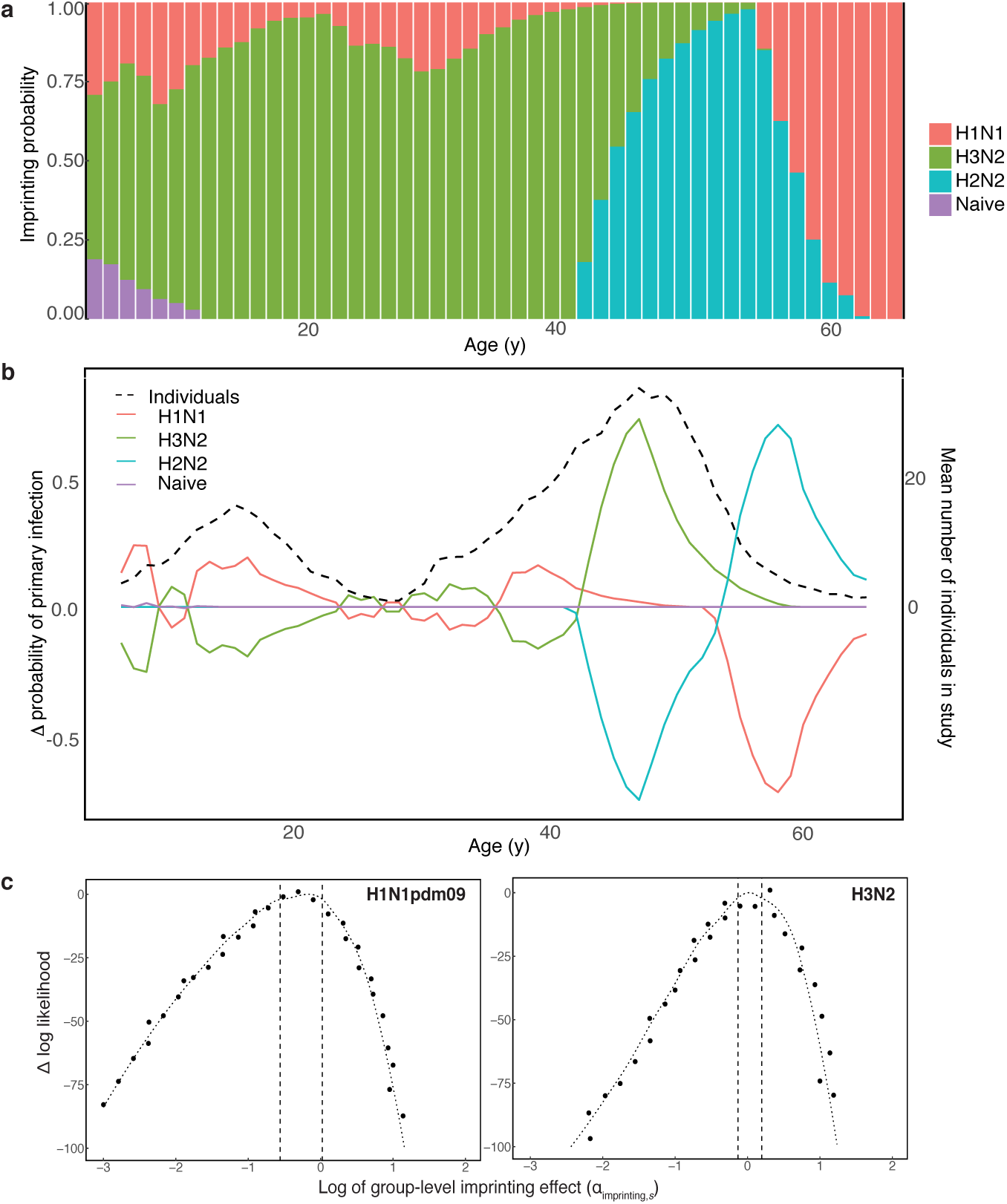
**(a)** Probability of imprinting by historically circulating influenza A subtypes by age in 2009. **(b)** Change in the mean probability of primary infection with historically-circulating subtypes by age between 2009 and 2014. The black dashed line gives the mean number of individuals by age that were observed in the data between 2009 and 2014. **(c)** Likelihood profiles for the effect of imprinting by H2N2 on the rate of infection with H1N1pdm09 (left) and the effect of imprinting by H3N2 on the rate of H3N2 infection (right). Values of the log parameter less than 0 (vertical dotted line) indicate a protective imprinting effect. The black dashed lines denote the 95% confidence interval.

**Figure S4:**
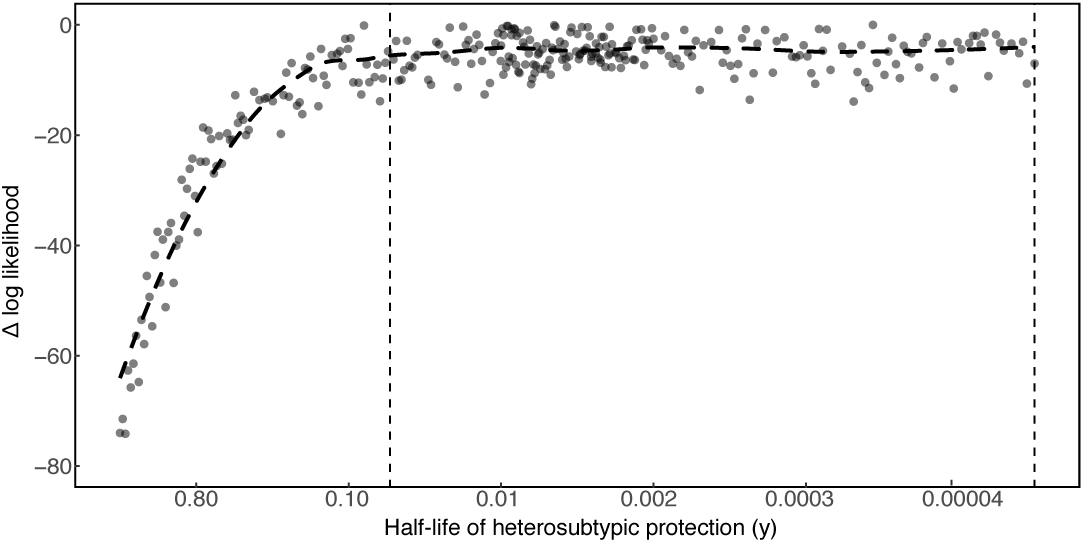
Likelihood profile of the rate of waning of heterosubtypic protection. The x-axis gives the corresponding half-life of protection.

**Figure S5:**
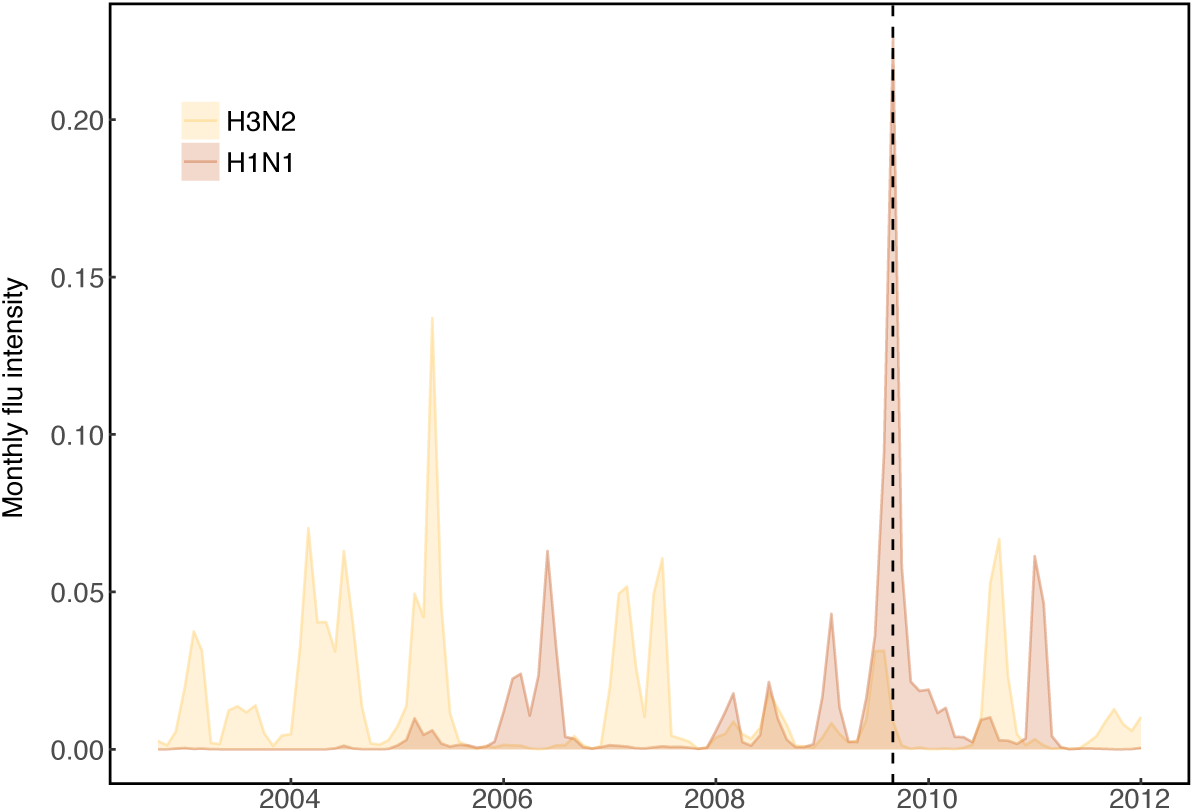
Subtype-specific flu intensity (ILI % positive) in Hong Kong. The black vertical dashed line denotes the earliest observation date in the data.

**Figure S6:**
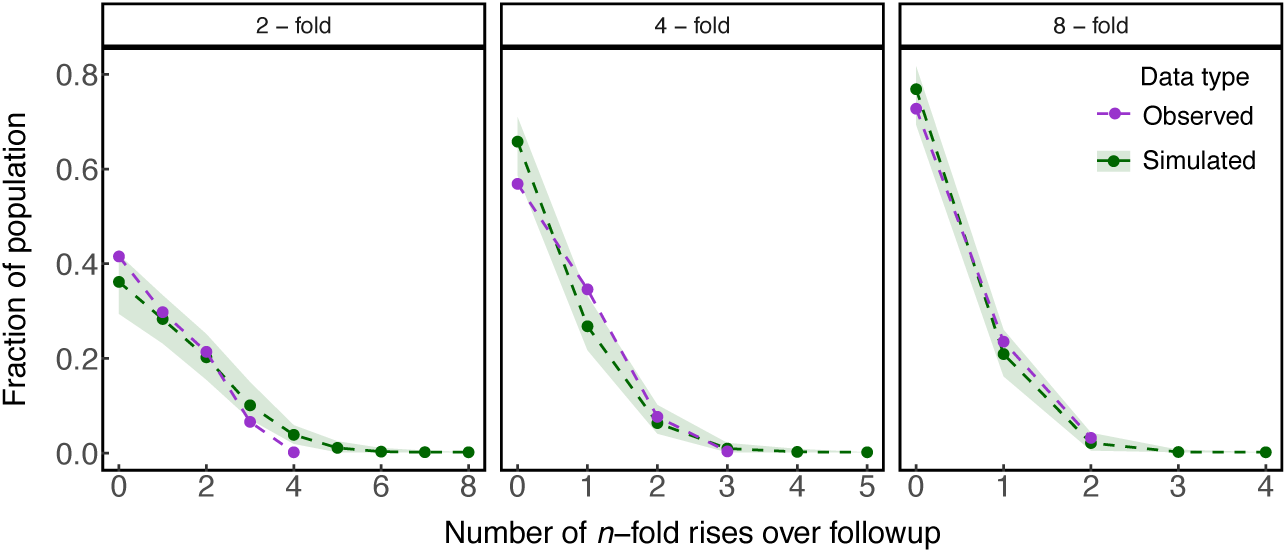
Observed and simulated distributions of consecutive 2-, 4-, and 8-fold titer rises per individual in the H1N1pdm09 data. The dashed blue lines give the medians from 1000 replicate simulations of the model, and the shaded blue areas are bounded by the 2.5% and 97.5% quantiles.

**Figure S7:**
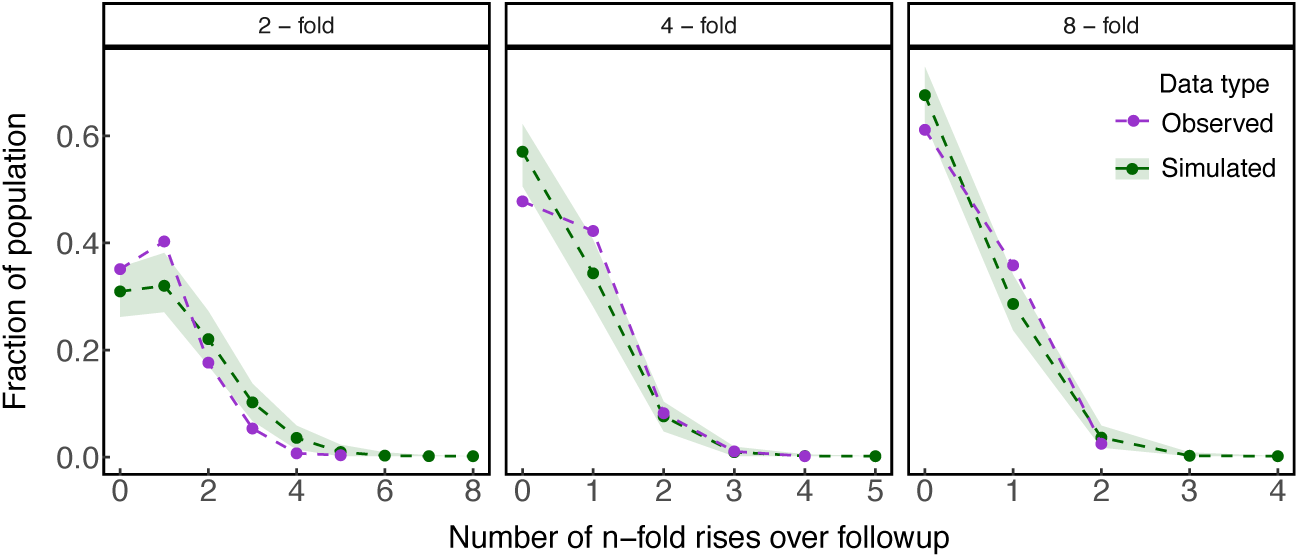
Observed and simulated distributions of consecutive 2-, 4-, and 8-fold titer rises per individual in the H3N2 data. The dashed blue lines give the medians from 1000 replicate simulations of the model, and the shaded blue areas are bounded by the 2.5% and 97.5% quantiles.

**Figure S8:**
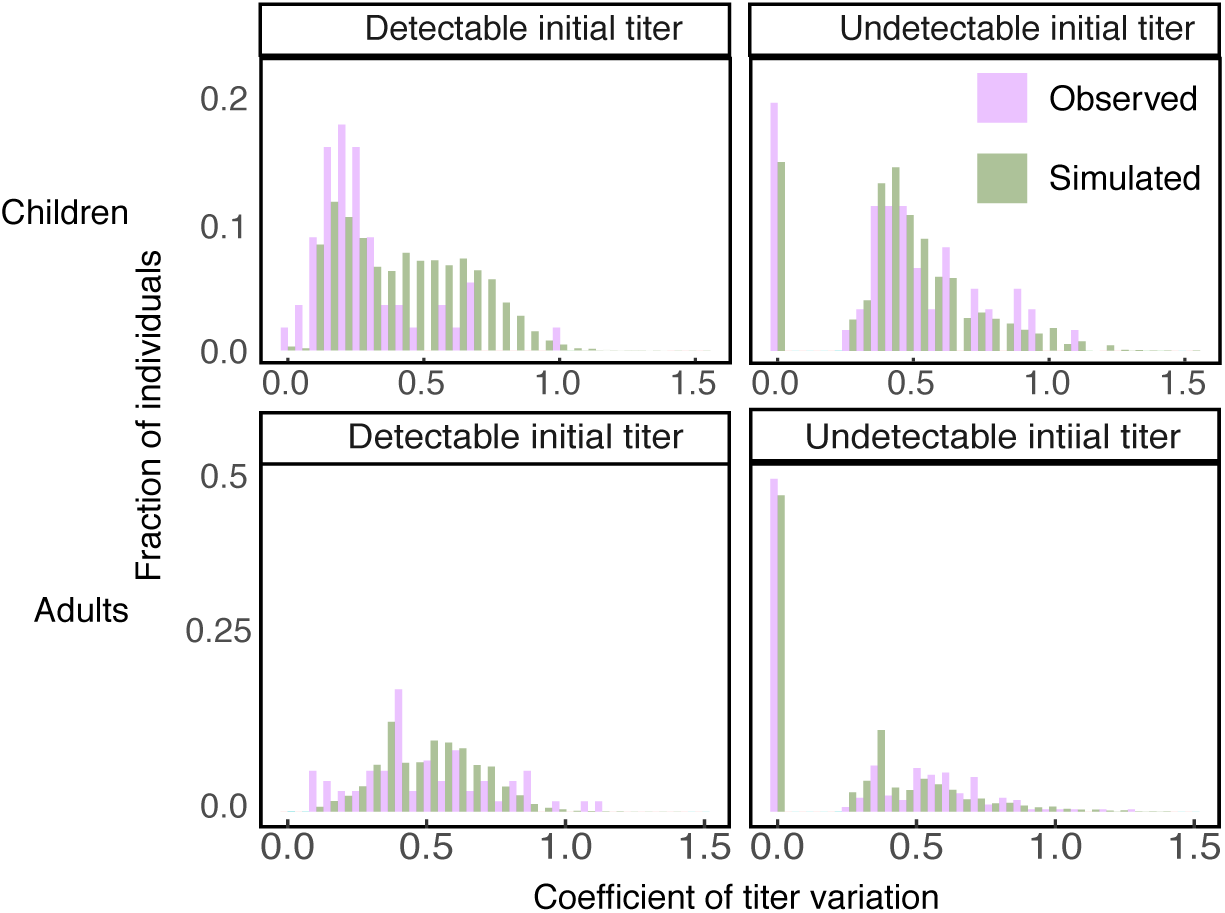
Observed and simulated distributions of the coefficients of log titer variation in the H1N1pdm09 data. The distributions are shown for individuals with initial titers *≥* 10 (detectable) and for individuals with initial titers <10 (undetectable).

**Figure S9:**
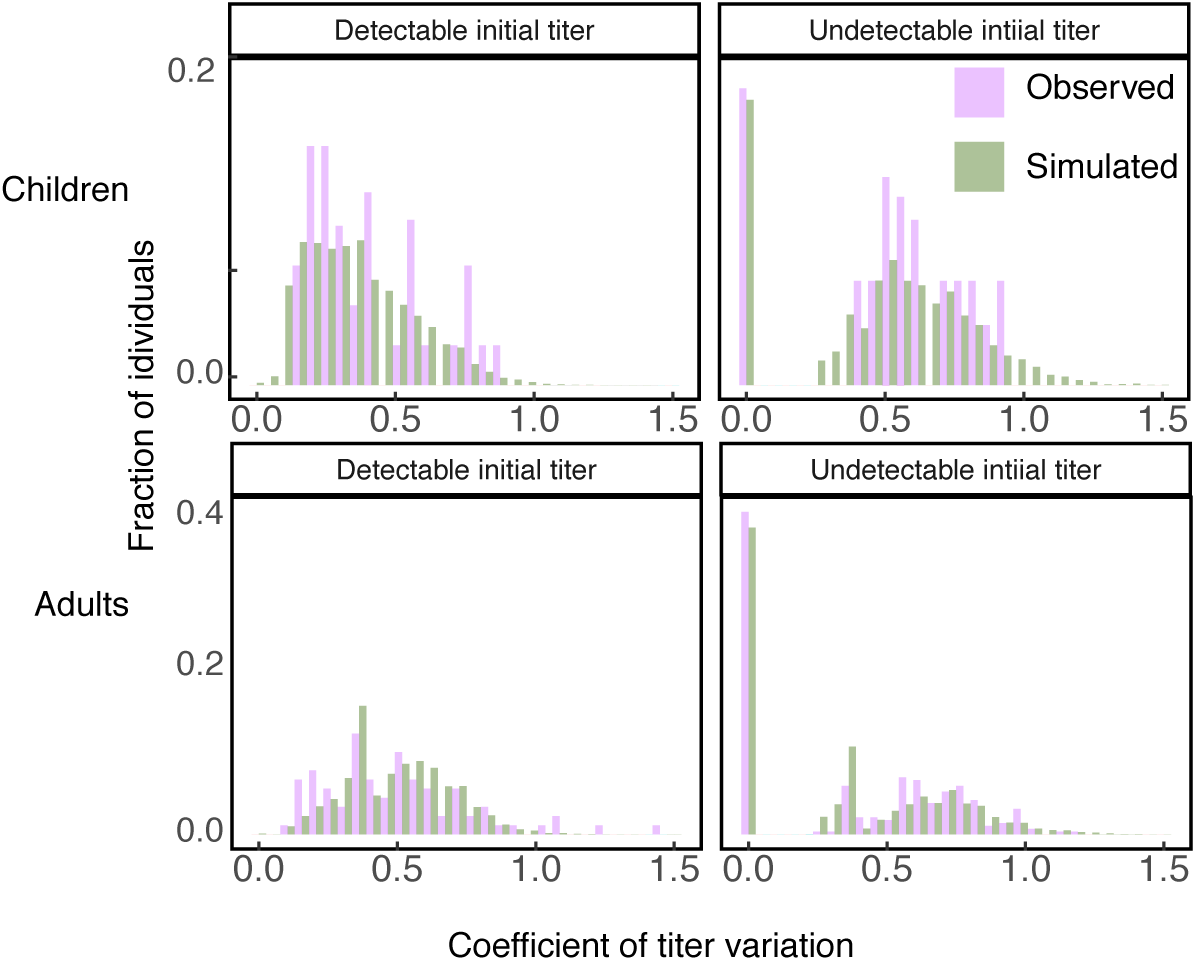
Observed and simulated distributions of coefficients of log titer variation in the H3N2 data. The distributions are shown for individuals with initial titers *≥* 10 (detectable) and for individuals with initial titers <10 (undetectable).

**Figure S10:**
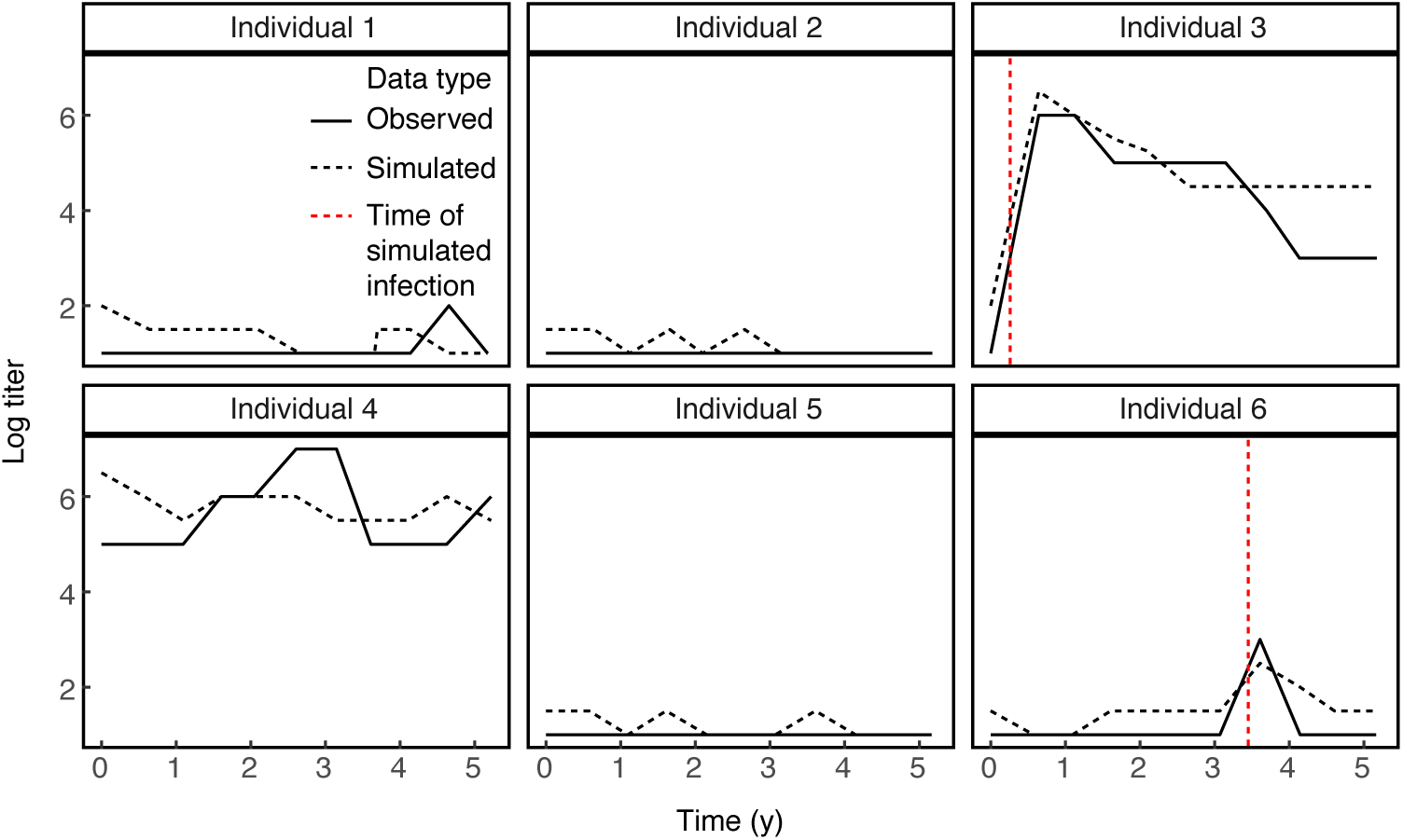
Simulated individual trajectories from the filtered particle population of the model for H1N1pdm09 at the maximum likelihood parameters. The filtered trajectories were obtained using 50,000 particles. The solid and dashed black lines give the observed log titer and the filtered log titer trajectory, respectively. The dashed red lines denote latent times of infection from the model. Results are shown for the first six individuals in the dataset.

**Figure S11:**
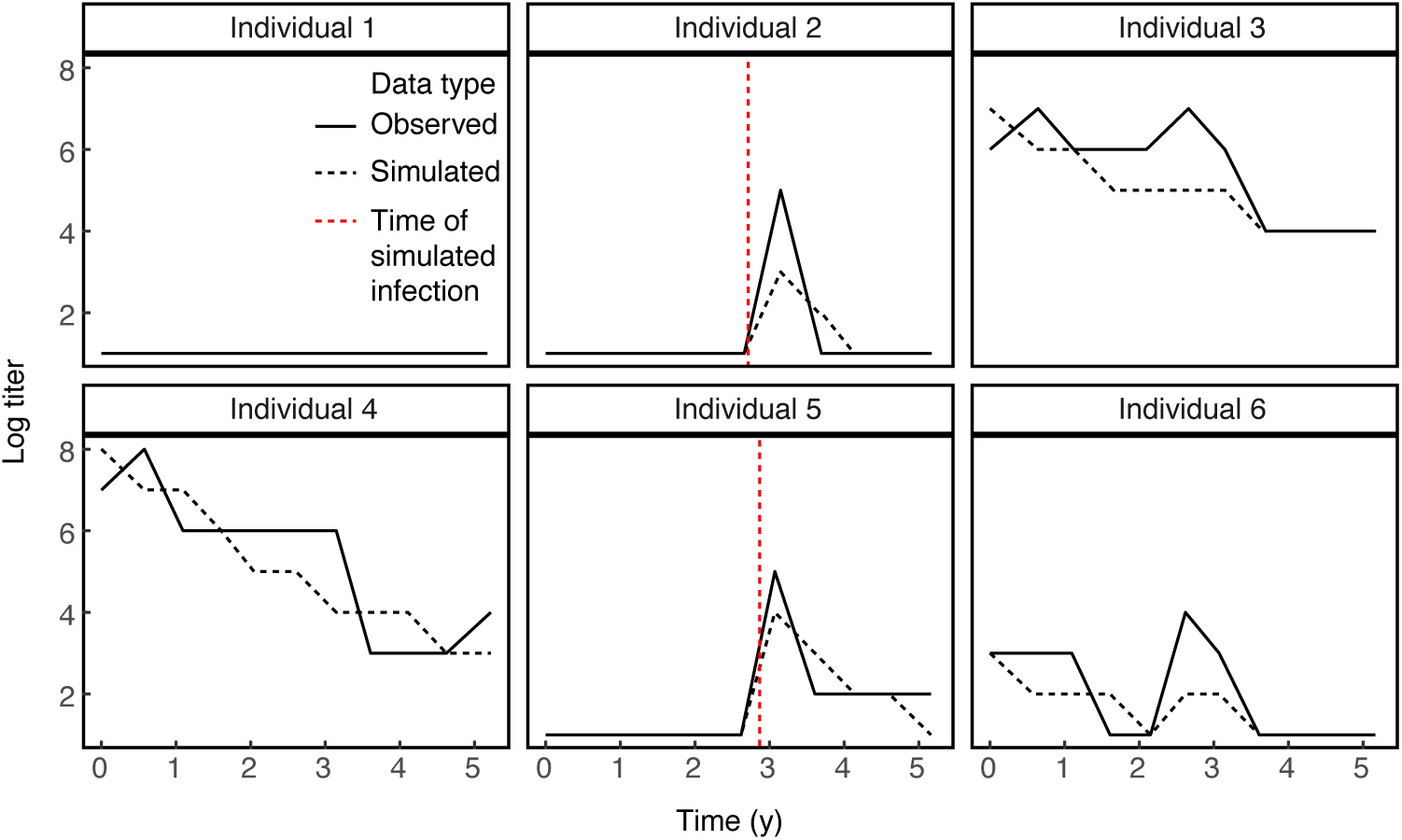
Simulated individual trajectories from the filtered particle population of the model for H3N2 at the maximum likelihood parameters. The filtered trajectories were obtained using 50,000 particles. The solid and dashed black lines give the observed log titer and the filtered log titer trajectory, respectively. The dashed red lines denote latent times of infection from the model. Results are shown for the first six individuals in the dataset.

**Figure S12:**
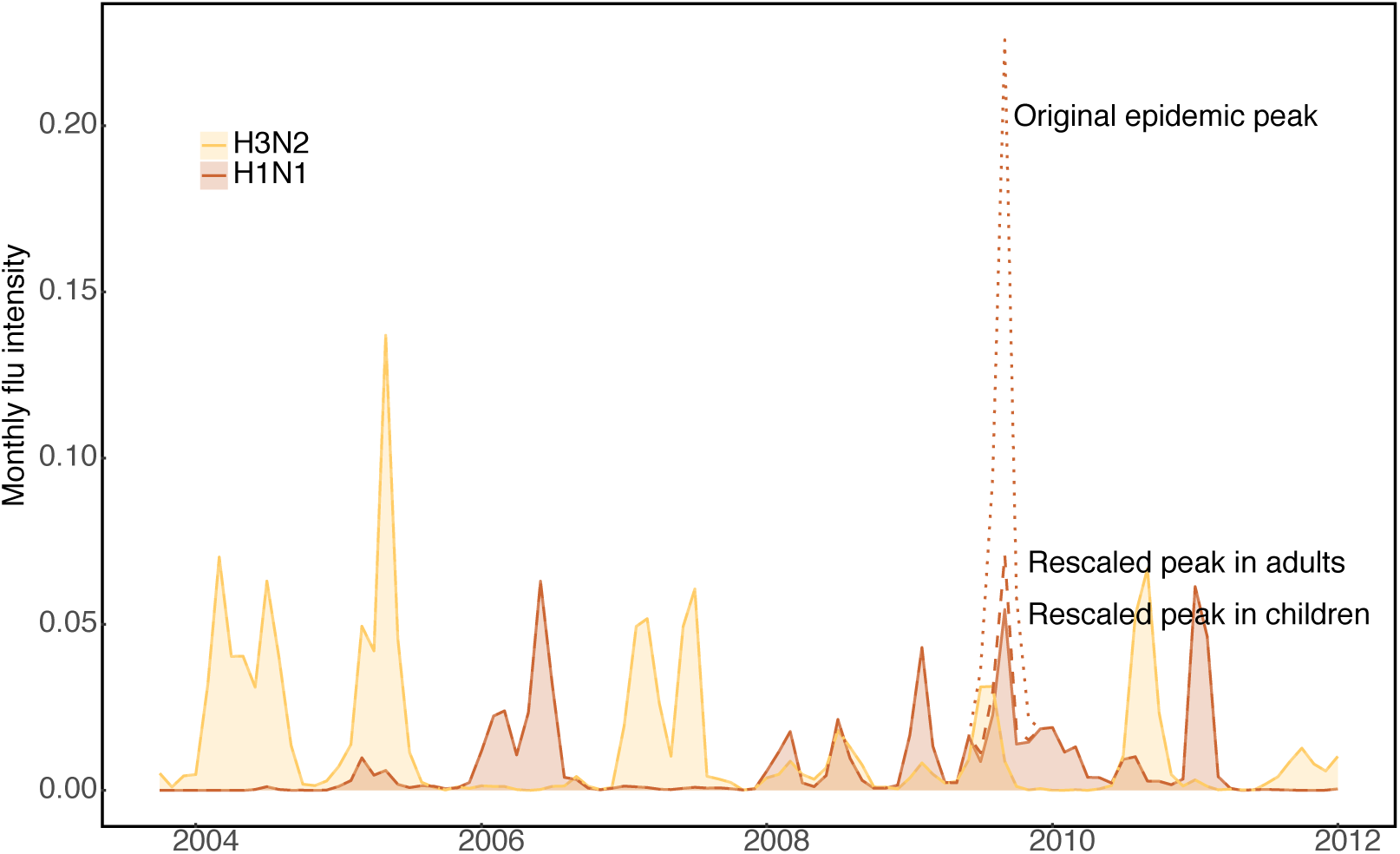
Rescaled community intensity of H1N1pdm09 during the 2009 pandemic in adults (dashed blue line) and in children (solid blue line and shading) compared to the original intensity reported by community surveillance (blue dotted line).

**Figure S13:**
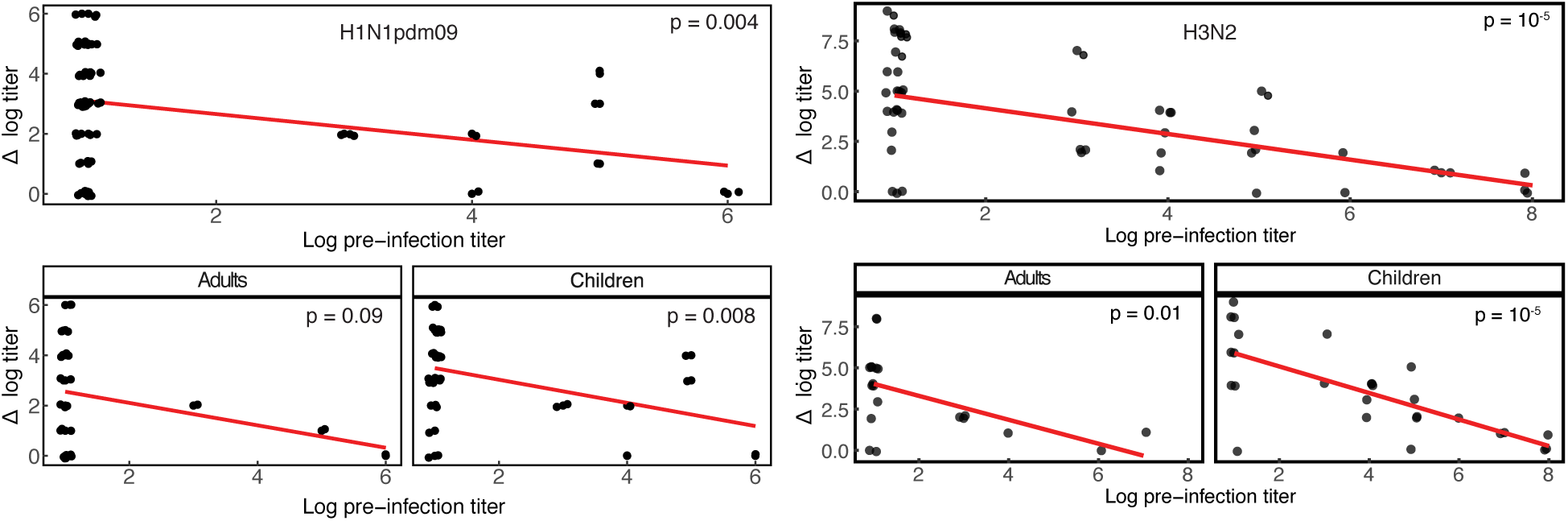
Observed titer boost as a function of the pre-infection log titer for individuals with PCR-confirmed infection with H1N1pdm09 (left) and H3N2 (right). Boosts are calculated as the post-infection minus the pre-infection log titer. The top panel for each subtype gives the relationship for aggregated data from children and adults. Note that log titers are defined as in Eq. 18.

**Figure S14:**
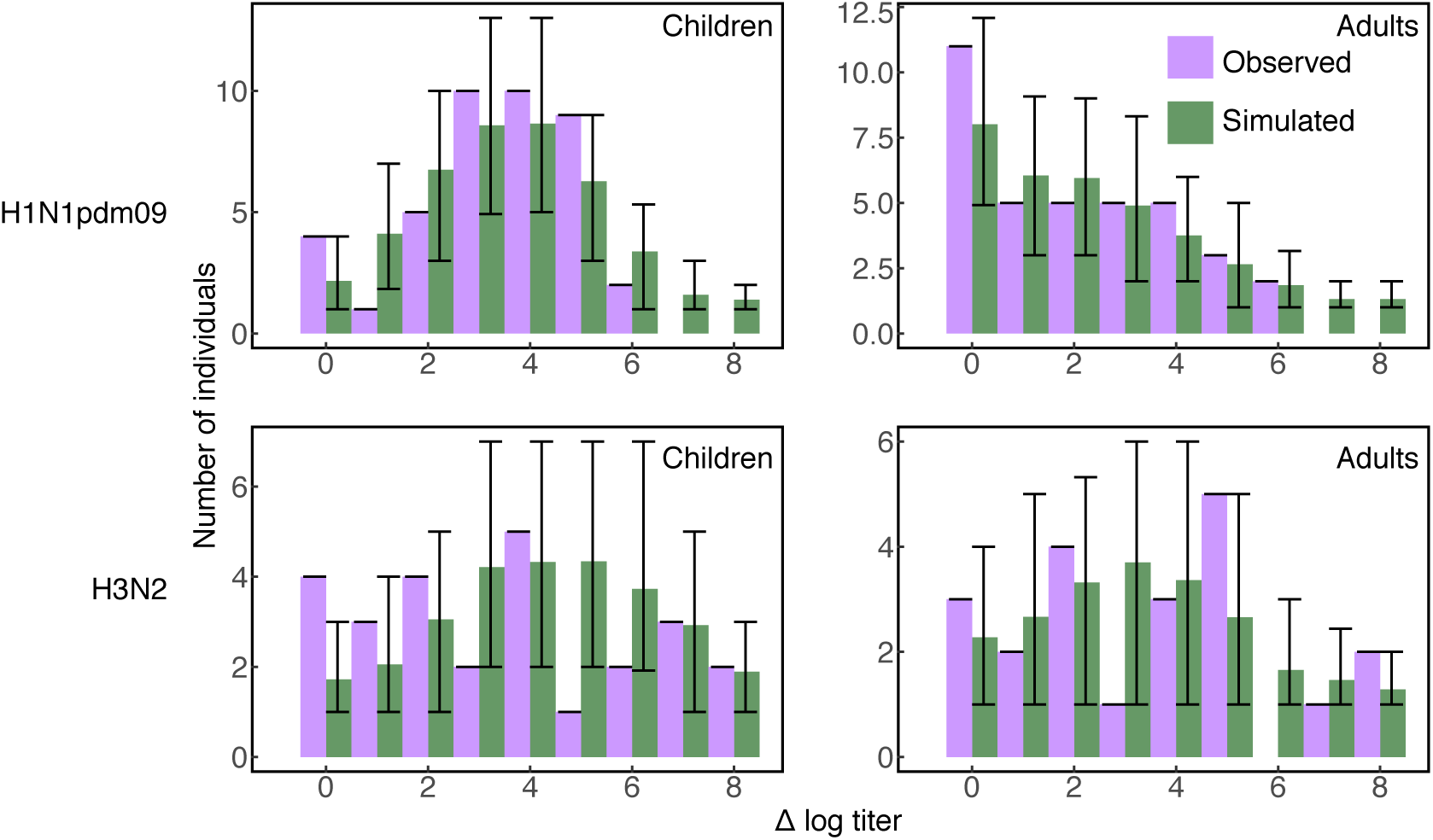
Observed and simulated distributions of titer boosts from the sub-model in individuals with PCR-confirmed infections. Boosts are calculated as the post-infection minus the pre-infection log titer. Error bars give the 95% CI among simulations.

**Figure S15:**
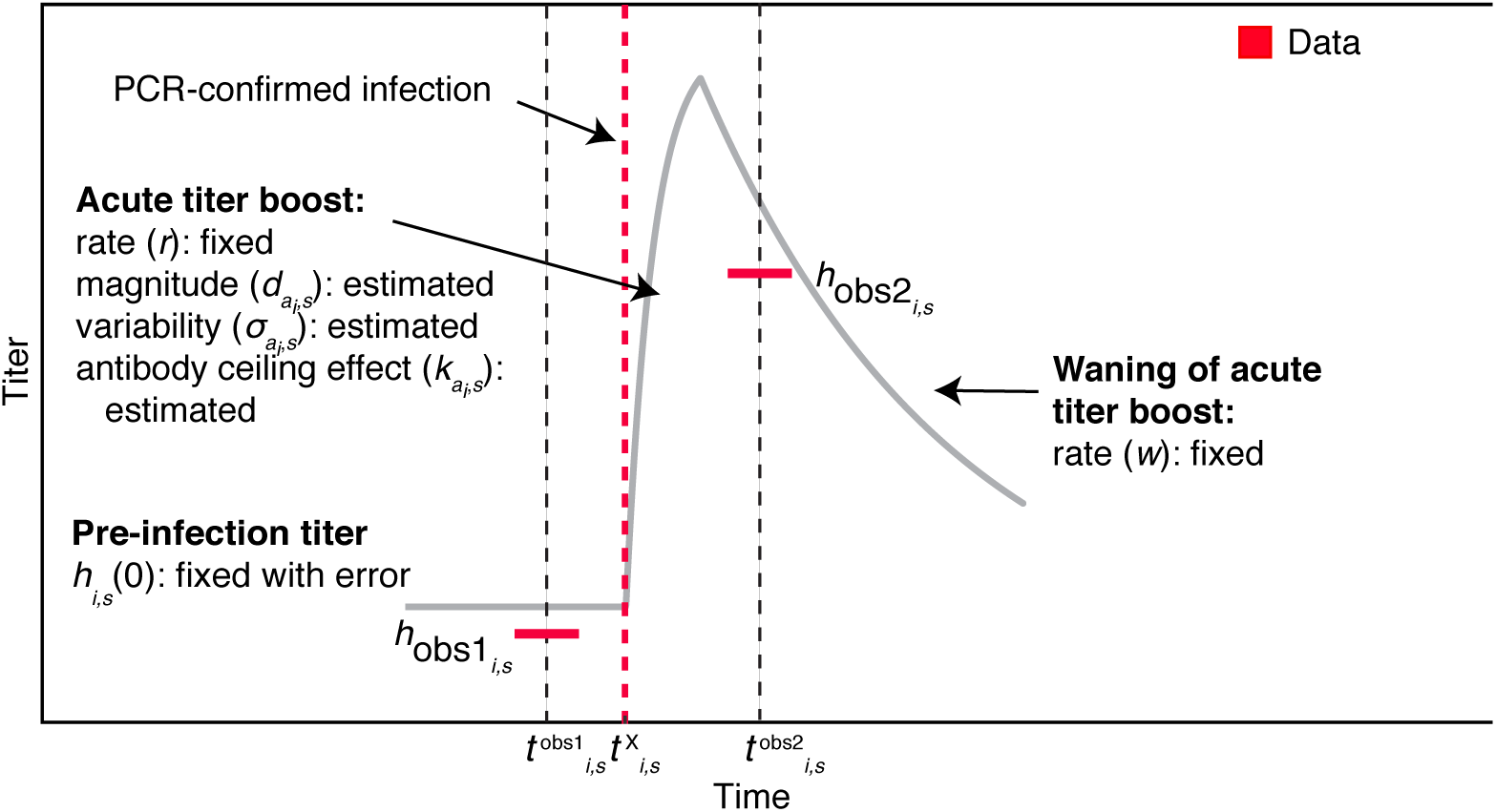
Schematic of the acute HI titer dynamics for individual *i* against subtype *s*, given infection at time. For the sub-model, is fixed based on the date of PCR-confirmed infection.

**Figure S16:**
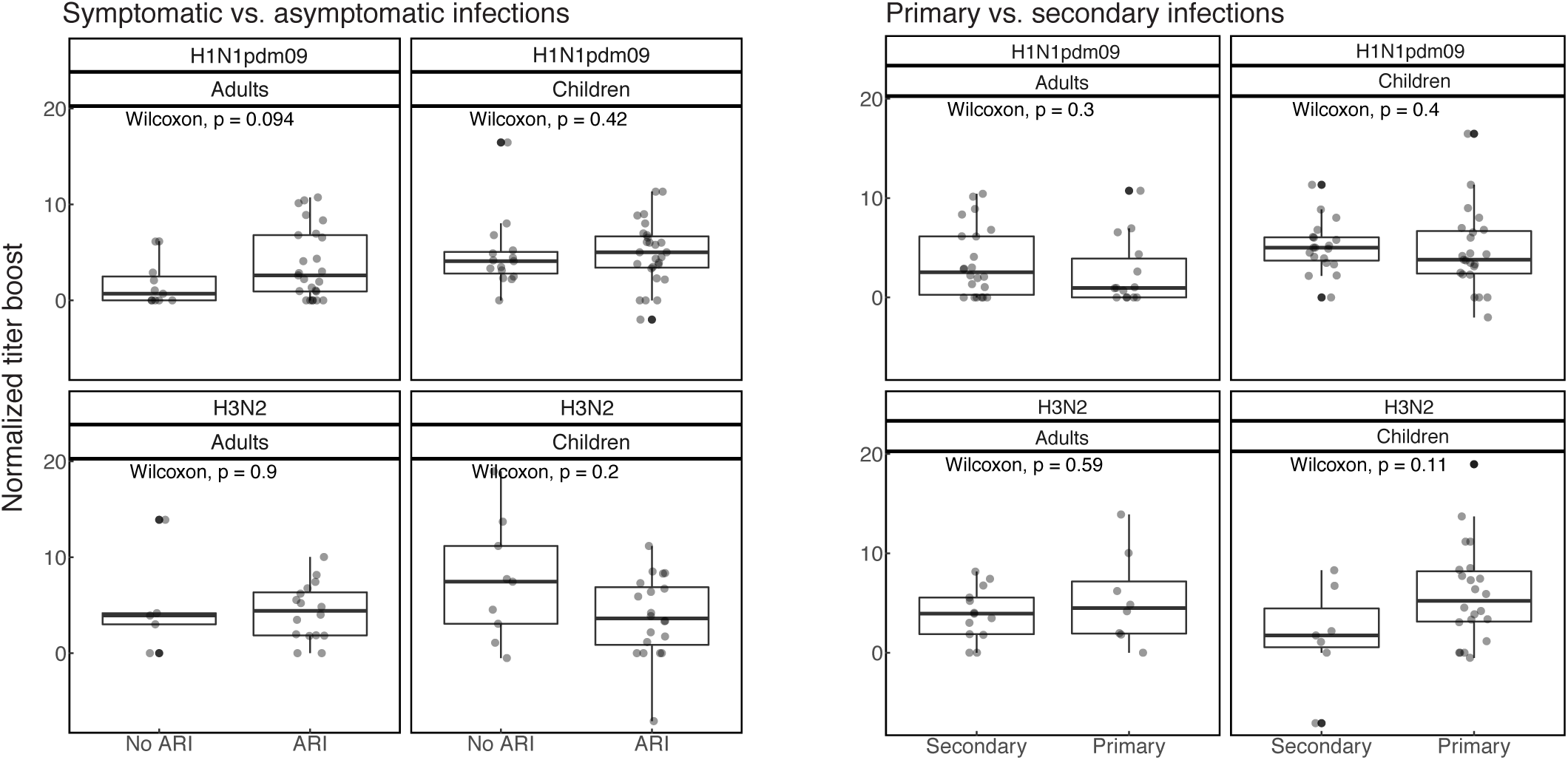
Distribution of normalized titer boosts after PCR-confirmed infections for symptomatic and asymptomatic infections (left) and primary and secondary infections (right). Normalized titer boosts are calculated as the log post-infection titer minus the log pre-infection titer divided by the length of time in years between the pre- and post-infection samples. Box plots give the median and interquartile range of the normalized titer boosts, and the individual data points are overlain with horizontal jitter. Differences in the mean of the distributions are determined by non-parametric Wilcoxon tests.

**Figure S17:**
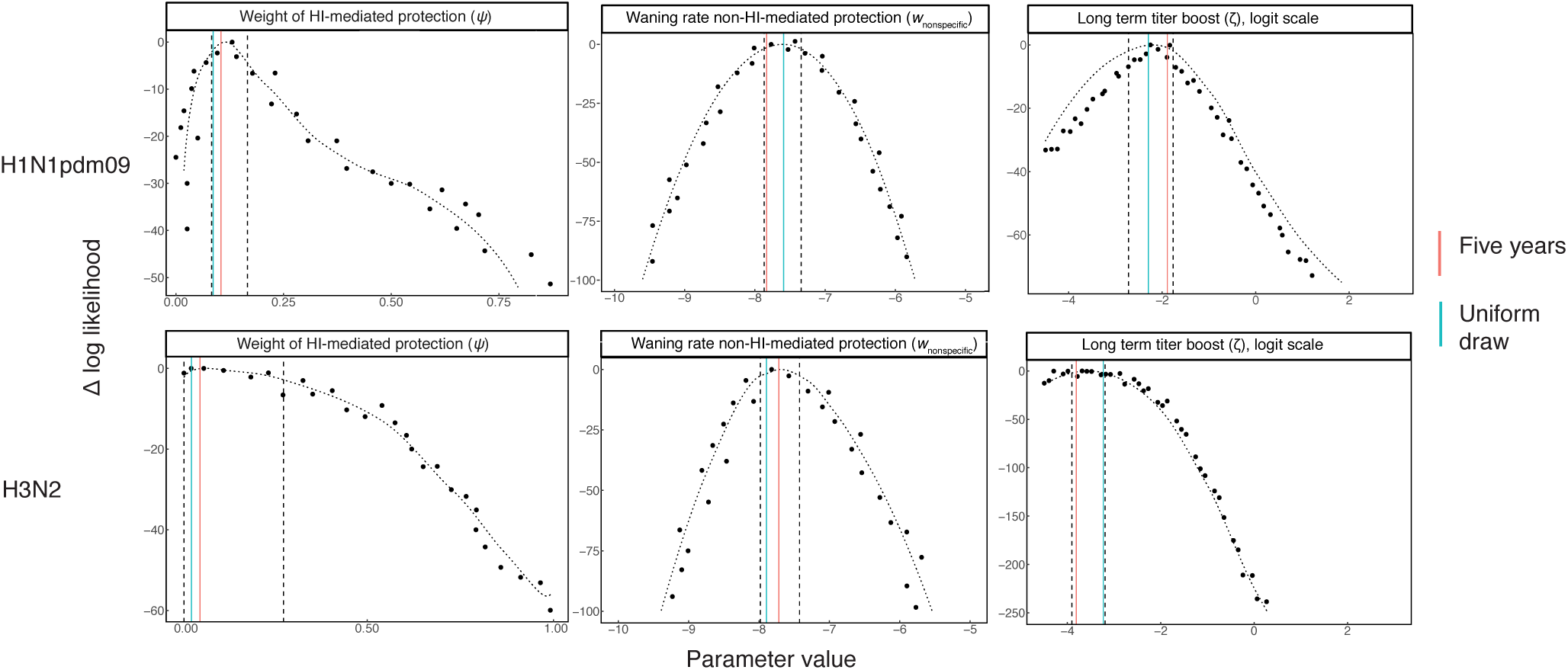
Likelihood profiles for the estimated parameters of the best-fit models in **adults** for H1N1pdm09 and H3N2. The dashed curve gives the spline computed by the Monte Carlo Adjusted Profile technique (Section S3). The vertical lines denote the MLEs from models under alternative initial conditions (section S2.3).

**Figure S18:**
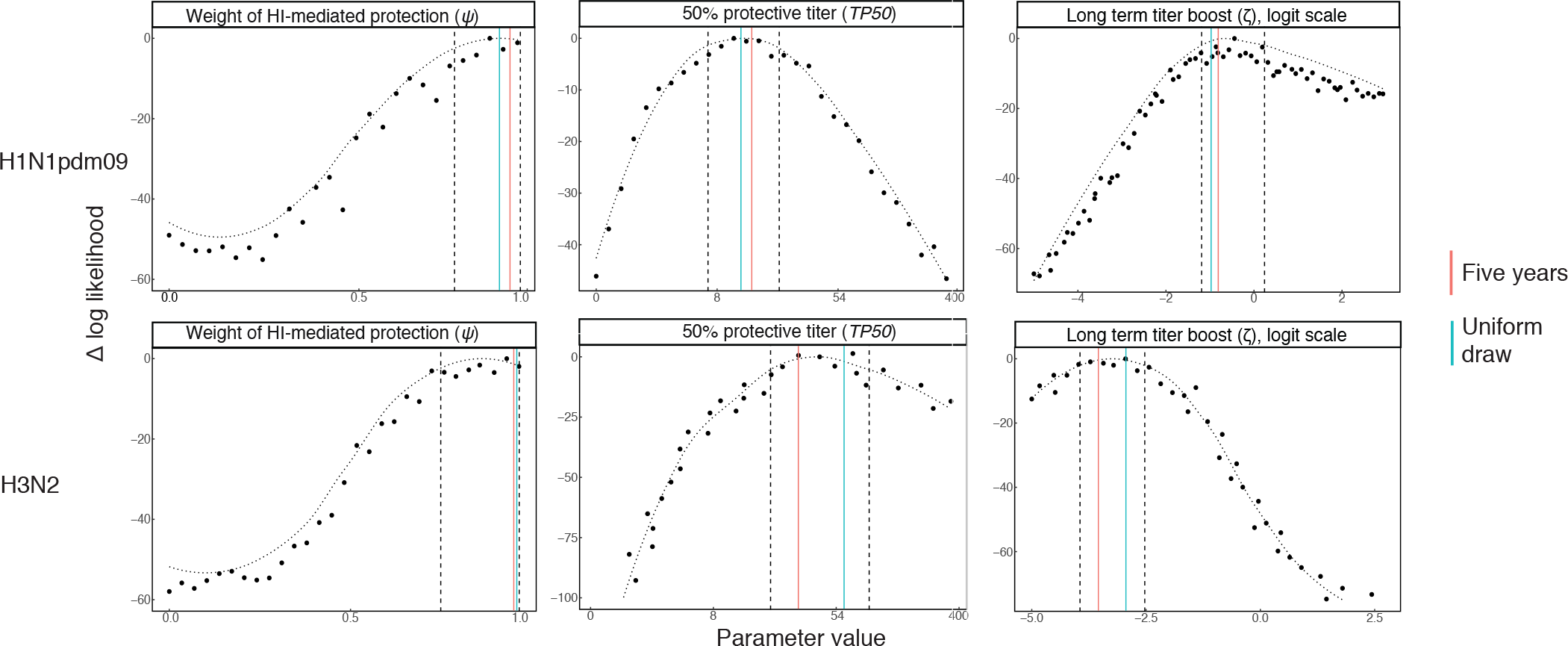
Likelihood profiles for the estimated parameters of the best-fit models in **children** for H1N1pdm09 and H3N2. The dashed curve gives the spline computed by the Monte Carlo Adjusted Profile technique (Section S3). The vertical lines denote the MLEs from models under alternative initial conditions (section S2.3).

**Figure S19:**
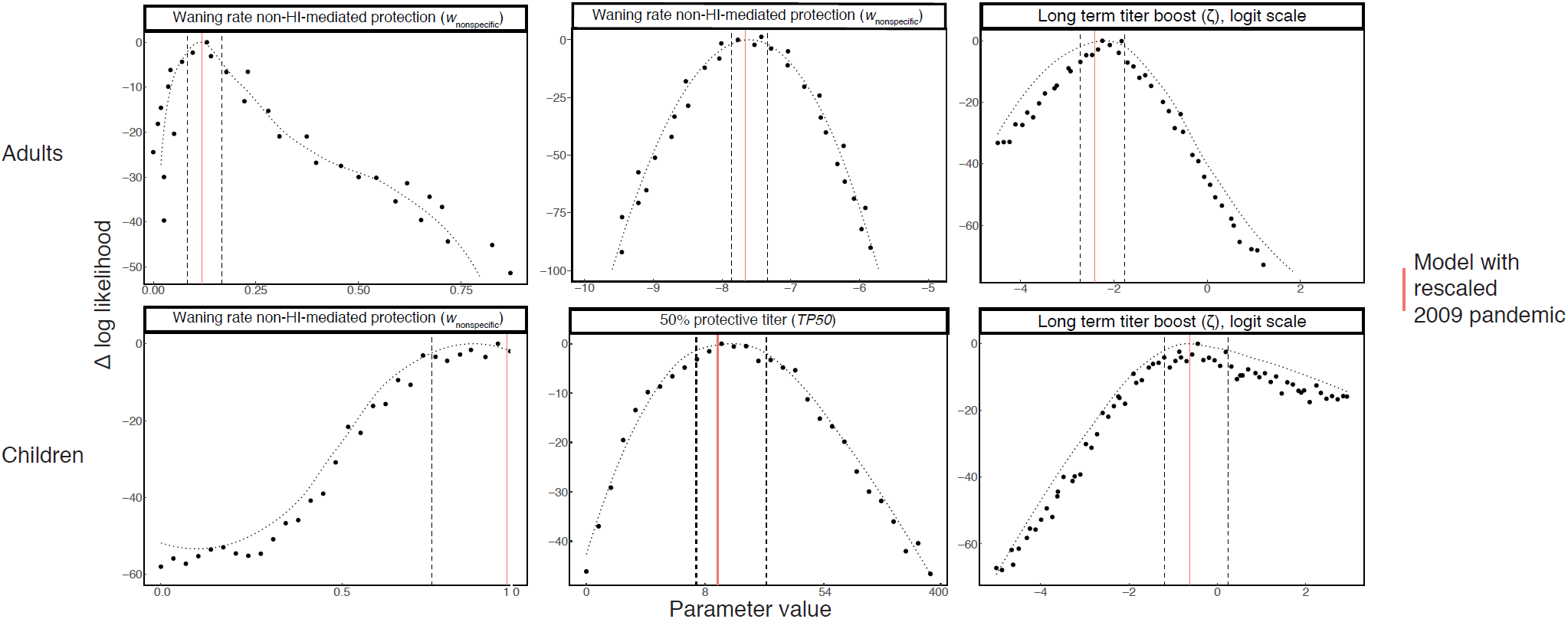
Likelihood profiles for the estimated parameters of the best-fit models in adults and children for H1N1pdm09. The dashed curve gives the spline computed by the Monte Carlo Adjusted Profile technique (section S3). The vertical lines denote the MLEs from the model with rescaled H1N1pdm09 intensity during the 2009 pandemic (section S2.4).

**Figure S20:**
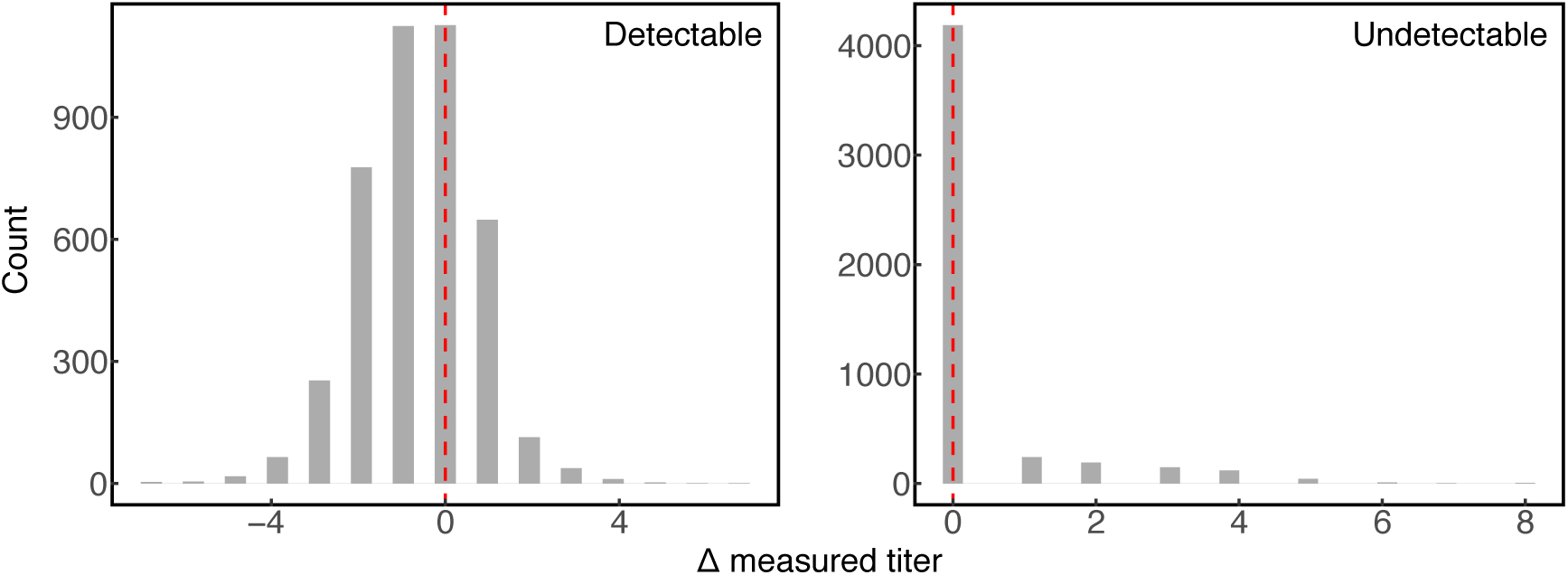
Distribution of difference between the second and first measured titer for sera that were tested twice, with distributions shown separately for detectable and undetectable titers based on the initial measurement. The vertical red line marks zero difference.

**Figure S21:**
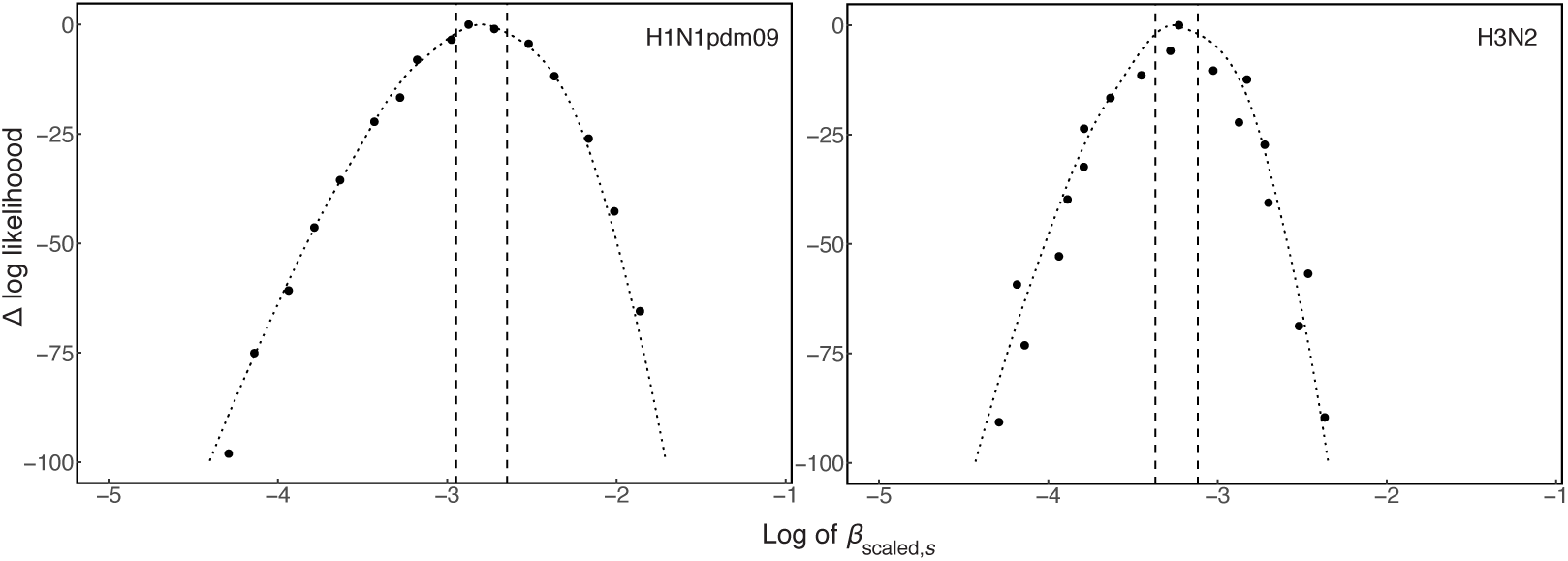
Likelihood profiles for the subtype-specific scaled transmission rate, *β*_scaled,*s*_for H1N1pdm09 and H3N2. The dashed horizontal line gives the threshold for statistical significance at a 95% level. The vertical lines denote the bounds of the 95% CI.

## 1 Short-term titer dynamics after PCR-confirmed infection

### 1.1 Model of short-term antibody boost after PCR-confirmed infection

To increase accuracy modeling the short-term post-infection titer dynamics (Eqs. 11-13, Fig. 1, Step 1a), we fit a “sub-model” to the observed titers before and after a PCR-positive swab (Fig. S15). We estimate the mean magnitude and variability of the short-term titer boost (*d*_*ai*,*s*_ and *σ*_*ai*,*s*_, respectively) and the dependence of the boost on the pre-infection titer, *k*_*ai*,*s*_. This allows us to test for the presence of an antibody ceiling effect, which has been identified in studies of post-vaccination titer dynamics [50, 59].

To fit the sub-models, we fixed the pre-infection latent titer, *h*_*i*,*s*_ (0), to the observed pre-swab titer, *h*_obs1,*i*,*s*_, allowing for two-fold uncertainty in the measured titer (Eq. 23). We fix the latent time of infection, 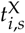, based on the date of the positive swab, assuming that the swab occurred during an infected period that we draw from a gamma distribution with fixed parameters (Table S4). We model the dynamics of the short-term titer rise as in Eq. 11, with the rate of rise *r* and time *T*_peak_ between infection and peak titer fixed (Table S4). After peaking, the titer wanes at rate *w* (fixed as in Table S4) until the time of the second observed value, *h*_obs2,*i*,*s*_.

### 1.2 Infection generates a variable short-term homosubtypic antibody boost that declines with increasing pre-infection titer

The raw data suggest an antibody ceiling effect (Fig. S13). We performed linear regressions of individuals’ observed changes in log titer on their observed pre-infection log titers, excluding one individual with ∆*t* > 1 y between the pre- and post-infection titer measurements. For both H1N1pdm09 and H3N2, the difference between pre- and post-infection log titers declines linearly with increasing pre-infection titer. The linear decline is statistically significant (p<0.02 for both subtypes). When we stratify the regressions in children and adults for each subtype, the decline is statistically significant, with p ≤0.01 for children for both subtypes and for adults with H3N2 (p=0.09 for adults with H1N1pdm09).

The dynamical sub-models also support an antibody ceiling effect for both subtypes in children and adults (Table S1), such that higher pre-infection titer diminishes the boost. For both subtypes, models that include the antibody ceiling effect (“with *k*”) outperform models that do not (“without *k*”, *k*_*ai*,*s*_ = 0) in children and adults (∆AICc > 2, Table S1). Therefore, part of the individual variation in the acute infection response can be explained by differences in pre-existing titers. Simulations from the MLEs of the best fit models of the short-term dynamics reproduce the shape of the observed distribution of titer boosts in children and adults after PCR-confirmed infection for both subtypes (Fig. S14).

From the maximum likelihood parameter estimates of the best-fit sub-models, we find substantial variability in antibody titer responses after PCR-confirmed infection with both subtypes in children and adults (Table S5). This finding is consistent with other analyses [51, 39]. The inferred standard deviation of the lognormal titer boost distribution (Eq. 14) ranges from 0.9 to 1.9 log titer units among children and adults for H1N1pdm09 and H3N2 (Table S5). The mean magnitude of the boost is higher for H3N2 than for H1N1pdm09 in both age groups. The variability in the acute infection response and the difference in the response between subtypes and age groups suggest that threshold titers used in sero-surveillance may not reliably predict infection in all individuals [10, 11].

### 1.3 Observed titer boosts secondary to symptomatic vs. asymptomatic infections and primary vs. secondary infections

The sub-model of the short-term titer dynamics does not distinguish between symptomatic infections and asymptomatic infections that may have been detected incidentally given illness in another household member. If the measured titer boosts vary with symptom severity, our estimates may be biased, since index cases were identified by symptoms. We define symptomatic infection by the presence of ARI in the two weeks before PCR-confirmed infection. Based on the household symptom diaries, approximately 70% of infections in both children and adults for both subtypes were symptomatic (Table S6). Children were more likely than adults for both subtypes to have a primary, or index case infection, meaning that no other household members had a PCR-confirmed infection or symptoms of an ARI in the two weeks prior to confirmed infection.

We compared the distributions of titer changes between symptomatic and asymptomatic infections and between primary and secondary infections (Fig. S16). Because titers wane, we normalized the boost by the interval between the pre- and post-infection sample dates. We find no statistically significant difference in the mean normalized titer boost between symptomatic and aymptomatic infections for either subtype in children or adults. Similarly, we find no statistically significant differences when comparing primary and secondary infections. Therefore, the data suggest that the titer boosts estimated from PCR-confirmed infections in adults and children are not biased by differences in asymptomatic case detection.

## 2 Model validation and sensitivity analysis

### 2.1 The model reproduces the observed distribution of titer rises among individuals

We compared the observed numbers of 2-,4-, and 8-fold increases in consecutive titer measurements for H3N2 and H1N1pdm09 to the distributions obtained from 1000 replicate simulations of the model at the MLEs (Figs. S7, S6). The model reproduces the observed distributions in children and adults for both subtypes.

### 2.2 The model overestimates the variation in individuals’ titers

We compared the observed distribution of the coefficient of titer variation for individuals to the distribution obtained from 1000 replicate simulations of the model at the MLE (Figs. S8, S9). We separately analyzed the distributions for individuals with detectable initial titers (≥ 10) and undetectable initial titers (<10). In our models, any simulated titer <10 takes the value 10 of an undetectable titer. Therefore, the variation in undetectable titers by measurement error alone is less than that for titers 10. For both subtypes, the models tend to overestimate the individual variation over time. The bias is more pronounced among individuals with detectable baseline titers, which might be explained by the measurement error. Nevertheless, the difference in the means of the observed and simulated CV distributions ranges from 0.0 to only 0.1 for children and adults with H1N1pdm09 and H3N2. Furthermore, the filtered particle population of the model, an estimate of the smoothed distribution of latent model variables, at the maximum likelihood parameters closely reproduces the observed titer trajectories for individuals (Figs. S10, S11).

### 2.3 The maximum likelihood parameter estimates are robust to assumptions about the initial conditions

To initialize the full model, we drew each individual’s time of most recent infection from the density of the subtype-specific influenza intensity in the seven years preceding the first observation. For comparison, we fitted the best-fit model for each subtype in children and adults using two alternative assumptions about the initial conditions. First, we drew the time of most recent infection from the density of the subtype-specific influenza intensity over the five years before the first observation (“Five years”, Figs. S17, S18). Second, we drew the time of most recent infection uniformly over the seven years before the first observation rather than using *L*_*s*_ (*t*) (“Uniform draw”, Figs. Figs. S17, S18). The maximum likelihood estimates of the alternative models fall within the 95% CI of the parameter estimates from the original assumption.

### 2.4 The inference results are robust to rescaling of the community intensity of H1N1pdm09 during the 2009 pandemic

During the first wave of pandemic influenza H1N1pdm09 in 2009, increased reporting rates and changes in health-care seeking behaviors affected surveillance [19, 60]. We re-fitted our models of H1N1pdm09 after scaling the community flu intensity to adjust for these differences. A previous study estimated separate scaling factors for the relationship between the H1N1pdm09 intensity proxy and the rate of infection before and after a November 2009 change point [61]. We rescaled our estimate *L*_*s*_ (*t*) of the 2009 pandemic H1N1pdm09 intensity by multiplying the intensity before the change point by the ratio *ρ* of the estimated post- and pre-change point scaling factors in children (*ρ* = 0.25) and adults (*ρ* = 0.29). Fig. S12 shows the rescaled intensity. Notably, our observations begin at the end of the 2009 pandemic. Fewer than 6% of observations in children and fewer than 5% of observations in adults occurred before the November 2009 change point. Fewer than 1% of observations in children and adults occurred before October 2009. The model recovered the same MLE given the rescaled pandemic intensity (Fig. S19).

### 2.5 The measurement error estimated from replicate titer measurements is consistent with literature estimates

The sera from three visit dates were measured twice. In our models, we used the first titer measurement for each serum sample (the measurement recorded closest to the sampling date). To estimate the measurement error, we calculated the difference in measured titer between the second and first replicates (Fig. S20). For detectable titer levels (>10), the standard deviation of the error distribution (SD = 1.23 log titer units) matches the measurement error that we fixed in the model according to estimates from the literature (Table S4). The negative central tendency of the difference between the second and first replicates among detectable titers (median = −0.98 log titer units log titer units) indicates that measured titer generally declines with time since sampling. Additionally, in line with previous analyses [15], we find that the error distribution is zero-inflated for undetectable titers <10 (Fig. S20), justifying our use of a separate measurement error for undetectable titers. A previous study estimated the probability of 2-fold measurement error for undetectable titers [15]. We therefore calculated the corresponding error (*c* = 0.74) in our normally distributed observation model that would yield the same probability of 2-fold measurement error for undetectable titers. The observation model is non-invertible (Eq. 19). Therefore, while we use the measurement error to draw simulated observations from a normal distribution centered around the latent log titers, we cannot back-calculate the value of the latent titers from observed data. This is why we assign the initial baseline titer *h*_baseline,*i*,*s*_ (0) from a possible two-fold range surrounding the lowest observed titer 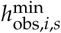 (Eq. 23).

### 2.6 Age-specific contact rates

We used age-specific contact rates estimated from a population-based survey of social contact patterns in Hong Kong that recorded daily contacts from over 1100 individuals in five age categories (Table S4, [57]). The authors calculated the relative number of contacts between individuals of each age category, adjusting for the propensity of individuals in each age class to respond to either paper or online questionnaires. The authors also reported the contact matrix between age groups. To fix the total daily number of contacts in our analysis for an individual of a particular age category, we multiplied the reported number of daily contacts from the reference group (children ages 0-10 years) by the relative number of contacts in each age category. To fix the number of daily contacts with individuals of each age group, we multiplied the total number of daily contacts by the fraction of age-group specific contacts from the contact matrix.

### 2.7 Historical influenza A subtype frequency data

Before 1968, annual subtype frequencies are specified by well-known durations of subtype circulation between historical pandemics [62]. After 1968, annual frequencies are calculated from subtype-specific surveillance data for Hong Kong or from Southeast Asia for years in which data from Hong Kong are unavailable. Between 1968 and 1997, subtype frequencies are the annual fraction of subtype-specific influenza A sequences in the Global Initiative on Sharing All Influenza Data (GISAID) database [63]. Aggregate regional data is used during years in which fewer than 30 sequences are available from Hong Kong or China. From 1997 to 2014, annual frequencies are the fraction of subtype-specific specimens reported by the Global Influenza Surveillance and Response System (GISRS) [64].

## 3 Monte Carlo error

A central feature of inference of stochastic models from large datasets is non-negligible Monte Carlo error that often makes it infeasible to calculate the likelihood with an error of less than one log likelihood unit. This principle holds especially for longitudinal (or panel) data, which often consist of a collection of individual time series that are dynamically independent apart from shared model parameters. Standard approaches for constructing 95% confidence intervals (CIs) rely on a threshold of 1.92 log likelihood units from the maximum log likelihood to construct the CI [65]. Therefore, high Monte Carlo error, or error in the likelihood calculation, also poses a challenge for estimating 95% confidence intervals. The Monte Carlo Adjusted Profile (MCAP) technique [53] shows that a quadratic approximation in the region of the maximum likelihood can be used to reliably extrapolate the 95% CI in systems with high Monte Carlo error. The MCAP algorithm accounts for the Monte Carlo error in the standard error of the spline fitted to a given likelihood profile. Importantly, Ionides and colleagues have shown that despite wider uncertainty around the maximum likelihood in systems with high Monte Carlo error, the MCAP approach reliably identifies the parameters. While the Monte Carlo variance of the log-likelihood estimate increases linearly with the amount of data, so too does the Fisher information, or the information about the system, and therefore the ability to reliably identify the maximum likelihood parameters.

In addition to using the MCAP technique to construct likelihood profiles, we accounted for Monte Carlo error in the maximum likelihood inference by initiating 100 independent MIF searches at random parameter values for any given model. To identify the MLE for a given point on a likelihood profile, we required that three MIF searches independently arrive within two log likelihood units of the maximum likelihood value.

## References

[1] D. C. Wiley, I. A. Wilson, and J. J. Skehel. Structural identification of the antibody-binding sites of Hong Kong influenza haemagglutinin and their involvement in antigenic variation. Nature, 289(5796):373–378, 1 1981.

[2] Andrew J. Caton, George G. Brownlee, Jonathan W. Yewdell, and Walter Gerhard. The antigenic structure of the influenza virus A/PR/8/34 hemagglutinin (H1 subtype). Cell, 31(2):417–427, 12 1982.

[3] Xiaocong Yu, Tshidi Tsibane, Patricia A McGraw, Frances S House, Christopher J Keefer, Mark D Hicar, Terrence M Tumpey, Claudia Pappas, Lucy A Perrone, Osvaldo Martinez, James Stevens, Ian A Wilson, Patricia V Aguilar, Eric L Altschuler, Christopher F Basler, James E Crowe, and Jr. Neutralizing antibodies derived from the B cells of 1918 influenza pandemic survivors. Nature, 455(7212):532–6, 9 2008.

[4] Kathy Hancock, Vic Veguilla, Xiuhua Lu, Weimin Zhong, Ebonee N. Butler, Hong Sun, Feng Liu, Libo Dong, Joshua R. DeVos, Paul M. Gargiullo, T. Lynnette Brammer, Nancy J. Cox, Terrence M. Tumpey, and Jacqueline M. Katz. Cross-Reactive Antibody Responses to the 2009 Pandemic H1N1 Influenza Virus. New England Journal of Medicine, 361(20):1945–1952, 11 2009.

[5] Sarah Cobey and Scott E Hensley. Immune history and influenza virus susceptibility. Current Opinion in Virology, 22:105–111, 2 2017.

[6] Fazekas de St Groth and R G Webster. Disquisitions of Original Antigenic Sin. I. Evidence in man. The Journal of Experimental Medicine, 124(3):331–45, 9 1966.

[7] Thomas Jr. Francis. On the Doctrine of Original Antigenic Sin. Proceedings of the American Philosophical Society, 104(6):572–578, 1960.

[8] Katelyn M Gostic, Monique Ambrose, Michael Worobey, and James O Lloyd-Smith. Potent protection against H5N1 and H7N9 influenza via childhood hemagglutinin imprinting. Science, 354(6313):722–726, 11 2016.

[9] Claudia Maria Trombetta and Emanuele Montomoli. Influenza immunology evaluation and correlates of protection: a focus on vaccines. Expert Review of Vaccines, 15(8):967–976, 8 2016.

[10] D Hobson, R L Curry, A S Beare, and A Ward-Gardner. The role of serum haemagglutination-inhibiting antibody in protection against challenge infection with influenza A2 and B viruses. The Journal of Hygiene (London), 70(4):767–77, 12 1972.

[11] Laurent Coudeville, Fabrice Bailleux, Benjamin Riche, FranÃğoise Megas, Philippe Andre, and Rene Ecochard. Relationship between haemagglutination-inhibiting antibody titres and clinical protection against influenza: development and application of a bayesian random-effects model. BMC Medical Research Methodology, 10(18):18, 12 2010.

[12] Hsiang-Yu Yuan, Marc Baguelin, Kin O. Kwok, Nimalan Arinaminpathy, Edwin van Leeuwen, and Steven Riley. The impact of stratified immunity on the transmission dynamics of influenza. Epidemics, 20(2017):84–93, 3 2017.

[13] Steven Black, Uwe Nicolay, Timo Vesikari, Markus Knuf, Giuseppe Del Giudice, Giovanni Della Cioppa, Theodore Tsai, Ralf Clemens, and Rino Rappuoli. Hemagglutination Inhibition Antibody Titers as a Correlate of Protection for Inactivated Influenza Vaccines in Children. The Pediatric Infectious Disease Journal, 30(12):1081–1085, 12 2011.

[14] Adam J. Kucharski, Justin Lessler, Jonathan M. Read, Huachen Zhu, Chao Qiang Jiang, Yi Guan, Derek A. T. Cummings, and Steven Riley. Estimating the Life Course of Influenza A(H3N2) Antibody Responses from Cross-Sectional Data. PLOS Biology, 13(3):e1002082, 3 2015.

[15] Simon Cauchemez, Peter Horby, Annette Fox, Le Quynh Mai, Le Thi Thanh, Pham Quang Thai, Le Nguyen Minh Hoa, Nguyen Tran Hien, and Neil M. Ferguson. Influenza Infection Rates, Measurement Errors and the Interpretation of Paired Serology. PLoS Pathogens, 8(12):e1003061, 12 2012.

[16] Xiahong Zhao, Yilin Ning, Mark Chen, and Alex R. Cook. Individual and population trajectories of influenza antibody titers over multiple seasons in tropical Singapore. American Journal of Epidemiology, 6 2017.

[17] Adam Kucharski, Justin Lessler, Derek Cummings, and Steven Riley. Timescales of influenza A/H3N2 antibody dynamics. bioRxiv, page 183111, 11 2017.

[18] Benjamin J Cowling, Kwok Hung Chan, Vicky J Fang, Lincoln L H Lau, Hau Chi So, Rita O P Fung, Edward S K Ma, Alfred S K Kwong, Chi-Wai Chan, Wendy W S Tsui, Ho-Yin Ngai, Daniel W S Chu, Paco W Y Lee, Ming-Chee Chiu, Gabriel M Leung, and Joseph S M Peiris. Comparative epidemiology of pandemic and seasonal influenza A in households. The New England Journal of Medicine, 362(23):2175–2184, 6 2010.

[19] B. J. Cowling, R. A. P. M. Perera, V. J. Fang, K.-H. Chan, W. Wai, H. C. So, D. K. W. Chu, J. Y. Wong, E. Y. Shiu, S. Ng, D. K. M. Ip, J. S. M. Peiris, and G. M. Leung. Incidence of Influenza Virus Infections in Children in Hong Kong in a 3-Year Randomized Placebo-Controlled Vaccine Study, 2009-2012. Clinical Infectious Diseases, 59(4):517–524, 8 2014.

[20] Annette Fox, Le Quynh Mai, Le Thi Thanh, Marcel Wolbers, Nguyen Le Khanh Hang, Pham Quang Thai, Nguyen Thi Thu Yen, Le Nguyen Minh Hoa, Juliet E. Bryant, Tran Nhu Duong, Dang Dinh Thoang, Ian G. Barr, Heiman Wertheim, Jeremy Farrar, Nguyen Tran Hien, and Peter Horby. Hemagglutination inhibiting antibodies and protection against seasonal and pandemic influenza infection. Journal of Infection, 70(2):187–196, 2 2015.

[21] Benjamin J Cowling, Wey Wen Lim, Ranawaka A P M Perera, Vicky J Fang, Gabriel M Leung, J S Malik Peiris, and Eric J Tchetgen Tchetgen. Influenza hemagglutination-inhibition antibody titer as a mediator of vaccine-induced protection for influenza B. Clinical infectious diseases: an official publication of the Infectious Diseases Society of America, 9 2018.

[22] Justin Lessler, Steven Riley, Jonathan M. Read, Shuying Wang, Huachen Zhu, Gavin J. D. Smith, Yi Guan, Chao Qiang Jiang, and Derek A. T. Cummings. Evidence for Antigenic Seniority in Influenza A (H3N2) Antibody Responses in Southern China. PLoS Pathogens, 8(7):e1002802, 7 2012.

[23] Raffael Nachbagauer, Angela Choi, Ruvim Izikson, Manon M. Cox, Peter Palese, and Florian Krammer. Age Dependence and Isotype Specificity of Influenza Virus Hemagglutinin Stalk-Reactive Antibodies in Humans. mBio, 7(1):01996–15, 2016.

[24] Yao-Qing Chen, Teddy John Wohlbold, Nai-Ying Zheng, Min Huang, Yunping Huang, Karlynn E. Neu, Jiwon Lee, Hongquan Wan, Karla Thatcher Rojas, Ericka Kirkpatrick, Carole Henry, Anna-Karin E. Palm, Christopher T. Stamper, Linda Yu-Ling Lan, David J. Topham, John Treanor, Jens Wrammert, Rafi Ahmed, Maryna C. Eichelberger, George Georgiou, Florian Krammer, and Patrick C. Wilson. Influenza Infection in Humans Induces Broadly Cross-Reactive and Protective Neuraminidase-Reactive Antibodies. Cell, 173(2):417–429, 4 2018.

[25] F M Davenport, Hennessy A V, T Francis, and P Fabisch. Epidemiologic and immunologic significance of age distribution of antibody to antigenic variants of influenza virus. The Journal of Experimental Medicine, 98(6):641–56, 12 1953.

[26] F. M. Davenport, A. V. Hennessy, C. H. Stuart-Harris, and T. Francis. Epidemiology of Influenza. Comparative Serological Observations in England and the United States. Lancet, pages 469–74, 1955.

[27] K M Sullivan, A S Monto, and I M Longini. Estimates of the US health impact of influenza. American journal of public health, 83(12):1712–6, 12 1993.

[28] Jerome I Tokars, Sonja J Olsen, and Carrie Reed. Seasonal Incidence of Symptomatic Influenza in the United States. Clinical Infectious Diseases, 66(10):1511–1518, 5 2018.

[29] Ira M. Longini, James S. Koopman, Arnold S. Monto, and John P. Fox. Estimating household and community transmission parameters for influenza. American Journal of Epidemiology, 115(5):736–751, 5 1982.

[30] Cecille Viboud, Pierre-Yves Boelle, Simon Cauchemez, Audrey Lavenu, Alain-Jacques Valleron, Antoine Flahault, and Fabrice Carrat. Risk factors of influenza transmission in households. The British journal of general practice: the journal of the Royal College of General Practitioners, 54(506):684–9, 9 2004.

[31] Frederick Hayden, Robert Belshe, Catalina Villanueva, Riin Lanno, Claire Hughes, Ian Small, Regina Dutkowski, Penelope Ward, and Jackie Carr. Management of Influenza in Households: A Prospective, Randomized Comparison of Oseltamivir Treatment With or Without Postexposure Prophylaxis. The Journal of Infectious Diseases, 189(3):440–449, 2 2004.

[32] Joshua G. Petrie, Suzanne E. Ohmit, Benjamin J. Cowling, Emileigh Johnson, Rachel T. Cross, Ryan E. Malosh, Mark G. Thompson, and Arnold S. Monto. Influenza Transmission in a Cohort of Households with Children: 2010-2011. PLoS ONE, 8(9):e75339, 9 2013.

[33] Rachel Savage, Michael Whelan, Ian Johnson, Elizabeth Rea, Marie LaFreniere, Laura C Rosella, Freda Lam, Tina Badiani, Anne-Luise Winter, Deborah J Carr, Crystal Frenette, Maureen Horn, Kathleen Dooling, Monali Varia, Anne-Marie Holt, Vidya Sunil, Catherine Grift, Eleanor Paget, Michael King, John Barbaro, and Natasha S Crowcroft. Assessing secondary attack rates among household contacts at the beginning of the influenza A (H1N1) pandemic in Ontario, Canada, April-June 2009: A prospective, observational study. BMC Public Health, 11(1):234, 12 2011.

[34] Jesse Papenburg, Mariana Baz, MarieâAŘÃĹve Hamelin, Chantal Rhéaume, Julie Carbonneau, Manale Ouakki, Isabelle Rouleau, Isabelle Hardy, Danuta Skowronski, Michel Roger, Hugues Charest, Gaston De Serres, and Guy Boivin. Household Transmission of the 2009 Pandemic A/H1N1 Influenza Virus: Elevated LaboratoryâĂŘ Confirmed Secondary Attack Rates and Evidence of Asymptomatic Infections. Clinical Infectious Diseases, 51(9):1033–1041, 11 2010.

[35] Alain Gagnon, Enrique Acosta, Stacey Hallman, Robert Bourbeau, Lisa Y Dillon, Nadine Ouellette, David J D Earn, D Ann Herring, Kris Inwood, Joaquin Madrenas, and Matthew S Miller. Pandemic Paradox: Early Life H2N2 Pandemic Influenza Infection Enhanced Susceptibility to Death during the 2009 H1N1 Pandemic. mBio, 9(1):02091–17, 1 2018.

[36] Edward Goldstein, Sarah Cobey, Saki Takahashi, Joel C. Miller, and Marc Lipsitch. Predicting the Epidemic Sizes of Influenza A/H1N1, A/H3N2, and B: A Statistical Method. PLoS Medicine, 8(7):e1001051, 7 2011.

[37] T. Sonoguchi, H. Naito, M. Hara, Y. Takeuchi, and H. Fukumi. Cross-Subtype Protection in Humans During Sequential, Overlapping, and/or Concurrent Epidemics Caused by H3N2 and H1N1 Influenza Viruses. Journal of Infectious Diseases, 151(1):81–88, 1 1985.

[38] M. L. B. Hillaire, S. E. van Trierum, J. H. C. M. Kreijtz, R. Bodewes, M. M. Geelhoed-Mieras, N. J. Nieuwkoop, R. A. M. Fouchier, T. Kuiken, A. D. M. E. Osterhaus, and G. F. Rimmelzwaan. Cross-protective immunity against influenza pH1N1 2009 viruses induced by seasonal influenza A (H3N2) virus is mediated by virus-specific T-cells. Journal of General Virology, 92(10):2339–2349, 10 2011.

[39] Judith M. Fonville, Pieter L. A. Fraaij, Gerrie de Mutsert, Samuel H. Wilks, Ruud van Beek, Ron A. M. Fouchier, and Guus F. Rimmelzwaan. Antigenic Maps of Influenza A(H3N2) Produced With Human Antisera Obtained After Primary Infection. Journal of Infectious Diseases, 213(1):31–38, 1 2016.

[40] Jae-Keun Park, Alison Han, Lindsay Czajkowski, Susan Reed, Rani Athota, Tyler Bristol, Luz Angela Rosas, Adriana Cervantes-Medina, Jeffery K Taubenberger, and Matthew J Memoli. Evaluation of Preexisting Anti-Hemagglutinin Stalk Antibody as a Correlate of Protection in a Healthy Volunteer Challenge with Influenza A/H1N1pdm Virus. mBio, 9(1):02284–17, 1 2018.

[41] Matthew S Miller, Thomas J Gardner, Florian Krammer, Lauren C Aguado, Domenico Tortorella, Christopher F Basler, and Peter Palese. Neutralizing antibodies against previously encountered influenza virus strains increase over time: a longitudinal analysis. Science Translational Medicine, 5(198):198ra107, 8 2013.

[42] Jacob Bock Axelsen, Rami Yaari, Bryan T Grenfell, and Lewi Stone. Multiannual forecasting of seasonal influenza dynamics reveals climatic and evolutionary drivers. PNAS, 111(26):9538–9542, 2014.

[43] Xiangjun Du, Aaron A King, Robert J Woods, and Mercedes Pascual. Evolution-informed forecasting of seasonal influenza A (H3N2). Science Translational Medicine, 9(413):eaan5325, 10 2017.

[44] Kristie M Grebe, Jonathan W Yewdell, and Jack R Bennink. Heterosubtypic immunity to influenza A virus: where do we stand? Microbes and Infection, 10(9):1024–9, 7 2008.

[45] J.H.C.M. Kreijtz, R.A.M. Fouchier, and G.F. Rimmelzwaan. Immune responses to influenza virus infection. Virus Research, 162(1-2):19–30, 12 2011.

[46] Benjamin J. Cowling, Sophia Ng, Edward S. K. Ma, Calvin K. Y. Cheng, Winnie Wai, Vicky J. Fang, Kwok-Hung Chan, Dennis K. M. Ip, Susan S. Chiu, J.S. Malik Peiris, and Gabriel M. Leung. Protective Efficacy of Seasonal Influenza Vaccination against Seasonal and Pandemic Influenza Virus Infection during 2009 in Hong Kong. Clinical Infectious Diseases, 51(12):1370–1379, 12 2010.

[47] Saverio Caini, Peter Spreeuwenberg, Gabriela F. Kusznierz, Juan Manuel Rudi, Rhonda Owen, Kate Pennington, Sonam Wangchuk, Sonam Gyeltshen, Walquiria Aparecida Ferreira de Almeida, ClÃąudio Maierovitch Pessanha Henriques, Richard Njouom, Marie-Astrid Vernet, Rodrigo A. Fasce, Winston Andrade, Hongjie Yu, Luzhao Feng, Juan Yang, Zhibin Peng, Jenny Lara, Alfredo Bruno, DomÃl’nica de Mora, Celina de Lozano, Maria Zambon, Richard Pebody, Leticia Castillo, Alexey W. Clara, Maria Luisa Matute, Herman Kosasih, Nurhayati, Simona Puzelli, Caterina Rizzo, Herve A. Kadjo, Coulibaly Daouda, Lyazzat Kiyanbekova, Akerke Ospanova, Joshua A. Mott, Gideon O. Emukule, Jean-Michel Heraud, Norosoa Harline Razanajatovo, Amal Barakat, Fatima el Falaki, Sue Q. Huang, Liza Lopez, Angel Balmaseda, Brechla Moreno, Ana Paula Rodrigues, Raquel Guiomar, Li Wei Ang, Vernon Jian Ming Lee, Marietjie Venter, Cheryl Cohen, Selim Badur, Meral A. Ciblak, Alla Mironenko, Olha Holubka, Joseph Bresee, Lynnette Brammer, Phuong Vu Mai Hoang, Mai Thi Quynh Le, Douglas Fleming, Clotilde El-Guerche Séblain, FranÃğois Schellevis, and John Paget. Distribution of influenza virus types by age using case-based global surveillance data from twenty-nine countries, 1999-2014. BMC Infectious Diseases, 18(1):269, 12 2018.

[48] Centre for Health Protection. Sentinel surveillance: Hong Kong Special Administrative Region, China, 2014.

[49] Andrew J. Dunning. A model for immunological correlates of protection. Statistics in Medicine, 25(9):1485–1497, 5 2006.

[50] Robert M Jacobson, Diane E Grill, Ann L Oberg, Pritish K Tosh, Inna G Ovsyannikova, and Gregory A Poland. Profiles of influenza A/H1N1 vaccine response using hemagglutination-inhibition titers Robert. Human Vaccines & Immunotherapeutics, 11(4):961–969, 4 2015.

[51] G Freeman, R A P M Perera, E Ngan, V J Fang, S Cauchemez, D K M Ip, J S M Peiris, and B J Cowling. Quantifying homologous and heterologous antibody titre rises after influenza virus infection. Epidemiology and infection, 144(11):2306–16, 8 2016.

[52] Edward L Ionides, C Bretó, and Aaron A King. Inference for nonlinear dynamical systems. Proceedings of the National Academy of Sciences, 103(49):18438–18443, 12 2006.

[53] E. L. Ionides, C. Breto, J. Park, R. A. Smith, and A. A. King. Monte Carlo profile confidence intervals for dynamic systems. Journal of The Royal Society Interface, 14(132):20170126, 2017.

[54] Clifford M. Hurvich and Chih-Ling Tsai. Regression and Time Series Model Selection in Small Samples. Biometrika, 76(2):297, 6 1989.

[55] R Bodewes, G de Mutsert, F R M van der Klis, M Ventresca, S Wilks, D J Smith, M Koopmans, R A M Fouchier, A D M E Osterhaus, and G F Rimmelzwaan. Prevalence of antibodies against seasonal influenza A and B viruses in children in Netherlands. Clinical and Vaccine Immunology, 18(3):469–76, 3 2011.

[56] W P Glezen, L H Taber, A L Frank, W C Gruber, and P A Piedra. Influenza virus infections in infants. The Pediatric infectious disease journal, 16(11):1065–8, 11 1997.

[57] Kathy Leung, Mark Jit, Eric H. Y. Lau, and Joseph T. Wu. Social contact patterns relevant to the spread of respiratory infectious diseases in Hong Kong. Scientific Reports, 7(1):7974, 2017.

[58] Anthony E Fiore, David K Shay, Karen Broder, John K Iskander, Timothy M Uyeki, Gina Mootrey, Joseph S Bresee, Nancy S Cox, Centers for Disease Control and Prevention (CDC), and Advisory Committee on Immunization Practices (ACIP). Prevention and control of influenza: recommendations of the Advisory Committee on Immunization Practices (ACIP), 2008. MMWR. Recommendations and reports: Morbidity and mortality weekly report. Recommendations and reports, 57(RR-7):1–60, 8 2008.

[59] SE Ohmit, JG Petrie, and RT Cross. Influenza hemagglutination-inhibition antibody titer as a correlate of vaccine-induced protection. The Journal of Infectious Diseases, 204(12):1879–1885, 2011.

[60] V. Marmara, A. Cook, and A. Kleczkowski. Estimation of force of infection based on different epidemiological proxies: 2009/2010 Influenza epidemic in Malta. Epidemics, 9:52–61, 12 2014.

[61] Tim K. Tsang, Simon Cauchemez, Ranawaka A. P. M. Perera, Guy Freeman, Vicky J. Fang, Dennis K. M. Ip, Gabriel M. Leung, Joseph Sriyal Malik Peiris, and Benjamin J. Cowling. Association Between Antibody Titers and Protection Against Influenza Virus Infection Within Households. The Journal of Infectious Diseases, 210(5):684–692, 9 2014.

[62] Edwin D Kilbourne. Influenza pandemics of the 20th century. Emerging infectious diseases, 12(1):9–14, 1 2006.

[63] Yuelong Shu and John McCauley. GISAID: Global initiative on sharing all influenza data - from vision to reality. Eurosurveillance, 22(13):30494, 3 2017.

[64] Antoine Flahault, Valentina Dias-Ferrao, Philippe Chaberty, Karin Esteves, Alain-Jacques Valleron, and Daniel Lavanchy. FluNet as a Tool for Global Monitoring of Influenza on the Web. JAMA, 280(15):1330, 10 1998.

[65] Ben Bolker. Likelihood and all that. In Ecological Models and Data in R, chapter 6, pages 169–221. Princeton University Press, 508 edition, 2007.

